# Chronobot: Deep learning guided time-resolved cryo-EM captures molecular choreography of RecA in homology search

**DOI:** 10.1101/2025.04.21.649820

**Authors:** Märt-Erik Mäeots, Simon Tupin, Mohammad M. N. Esfahani, Juan B. Rodriguez Molina, Julie A. Clapperton, Aran Amin, Albane Imbert, Radoslav I. Enchev

**Affiliations:** Visual Biochemistry Laboratory, The Francis Crick Institute, 1 Midland Rd, London NW1 1AT, United Kingdom; Making STP, The Francis Crick Institute, 1 Midland Rd, London NW1 1AT, United Kingdom; Imperial Advanced Hackspace, Enterprise, Imperial College London, London, W12 7TA, United Kingdom; Faculty of Life Science and Medicine, King’s College London, St Thomas’ Hospital, London, SE1 7EH, United Kingdom

## Abstract

The function of proteins and other biological macromolecules is regulated by conformational dynamics^1^. Many functional changes take place on millisecond timescales which cannot be experimentally captured by manual sample preparation for cryo-EM^2^. Here we introduce Chronobot, a robust, data-driven platform enabling reproducible, time-resolved cryo-EM sample preparation to visualize these transient intermediates. We quantify ice thickness, an important precondition and thus a reliable predictor of 3D reconstruction quality, using two complementary methods: a bespoke deep learning model analysing high-speed camera videos of the grid just before vitrification, and detailed ice thickness quantifications of outputs from common TEM screening workflows like the EPU software. Combining these methods enables rapid optimisation, resulting in an 11-fold improvement in cryo-EM sample quality compared to our previously reported workflow^3^.

To demonstrate the Chronobot in capturing transient reaction intermediates visualised through cryo-EM and single particle analysis we focused on RecA homology search. The RecA family of recombinases perform the essential task of rapidly scanning for homologous dsDNA sequences to initiate homologous recombination. The dynamics of these RecA-dsDNA interactions occur on millisecond timescales, limiting structural insights^4^. We capture time-resolved homology search intermediates at 250 milliseconds. These structures reveal the involvement of the secondary DNA binding site in initial capture of dsDNA before homology sampling occurs. We also observe three-strand homology sampling intermediates, where the homologous strand is not fully displaced, and homology is not stably bound. Our results suggest a model of how RecA-family recombinases function in early homologous recombination, by coordinating the incoming DNA between RecA’s various DNA binding sites depending on the stage of homology search and the presence of sufficient homology. We anticipate the Chronobot method to be broadly applicable to processes which cannot be captured by manual sample preparation methods. In addition, by leveraging AI inference, our rapid user feedback mechanisms allow for per-sample optimisation of grid conditions, increasing the likelihood of success and reducing the sample requirements of each time-resolved experiment.

## Introduction

Biological macromolecules such as proteins adopt conformational ensembles determined by the system’s free energy ^1^. Many protein fluctuations at equilibrium occur on microsecond timescales and are thermally driven, often unrelated to function. Functional conformational changes, however, involve specific subsets of states triggered by interactions, typically proceeding via transient intermediates over milliseconds to seconds, resulting in folding, allosteric regulation, substrate binding and/or catalysis^5^.

Visualizing transient conformational states is challenging because these states have low or no occupancy at steady state^5^. Manual sample preparation is significantly slower than the millisecond timescales critical for studying biochemical reactions. Among structural biology techniques — NMR, X-ray crystallography, and cryo-electron microscopy — cryo-EM has become popular for time-resolved studies due to recent technical advancements and broad applicability ^2,3,6–9^. Vitrification largely preserves the structural ensemble of macromolecules^10,11^, enabling redesigned sample preparation methods compatible with millisecond resolution and established data workflows.

Developing time-resolved sample preparation devices aims to reduce sample preparation times dramatically compared to manual methods, requiring solutions to technical challenges like rapid protein mixing^12^. Additionally, the sample must be quickly deposited as a thin (<100 nm) liquid film onto grids for vitrification, posing challenges for consistency, reproducibility, and quality of the vitreous thin sample film (hereafter referred to as “ice”) compared to slower blotting methods^13,14^. This limitation reduces reproducibility, increases sample demands, and affects the overall quality of the ice in many current methods. It is therefore imperative that continued progress is made in making these devices accommodate a broad timescale of reactions while producing high-quality data. Here we introduce the Chronobot, a next-generation time-resolved sample preparation device designed for robustness and reproducibility while providing the user fine-grained optimisation controls to tune grid preparation parameters to suit their sample. This is also supported by an array of sensors and feedback mechanisms to ensure accurate time resolution.

Improving ice thickness and uniformity is critical^15^ but challenging to measure in order to guide optimisation. Current methods offer high accuracy but require specialized imaging techniques (high magnification, tomography, vacuum reference images) ^16–19^. Ideal methods from a user perspective would estimate ice thickness without extra data collection during standard screening, or even outside the transmission electron microscope (TEM), enabling rapid iteration and discarding of low-quality grids. In this work, we present two new methods for automated ice thickness estimation. Our first method uses a U-net-derived^20^ algorithm trained on high-speed camera and TEM image pairs to predict ice thickness during grid plunging. Our second approach is based on the MeasureIce approach^19^. However, instead of using intermediate magnification, we attempt to quantify ice using the 150x magnification images collected by the EPU software as part of its standard screening workflow. We also eliminate the need to collect separate reference images by estimating these values from the unlabelled dataset. We demonstrate that while both methods result in reduced accuracy compared to rigorous methods, their ease of use as an integral part of the Chronobot workflow has significant practical value. We use these methods to pre-screen grids before the TEM to save screening time, aiding the user to obtain suitable samples in less time.

Time-resolved cryo-EM sample preparation workflow is uniquely positioned to address key outstanding biological questions with broad importance to basic biology and translational science. The homologous recombination machinery exemplifies such a system. Homologous recombination (HR) is one of the fundamental processes of life, not restricted to DNA repair. In bacteria, recombination is a key driver of genetic diversity and spreading of mutations^21^. In eukaryotes HR also provides an alternative pathway for telomere maintenance^22^, and underlies genetic reshuffling during meiosis^23^. Central to HR is the homology search, orchestrated by RecA-family recombinases (e.g., RecA, Rad51), where DNA is opened, homology is identified, and suitable sequences are captured prior to strand exchange^4,24,25^.

The homology search begins with RecA polymerizing onto single-stranded DNA (ssDNA), holding the ssDNA in site I^26^, forming a presynaptic filament that binds and locally destabilizes double-stranded DNA (dsDNA), sampling for complementarity in the ensuing dsDNA bubble region^27–29^. Single molecule imaging revealed that both RecA and Rad51 require 8-nt of consecutive homology to capture dsDNA for more than a few hundred milliseconds and eventually form a stable D-loop^4^.

Molecular dynamics simulations have hinted at rapid exchange of base pairs between the three strands, aided by the L2 loop and Site II of RecA which counteract reannealing of the dsDNA bubble^27^. Single-molecule fluorescence measurements have shown the rapid rejection of unsuitable sequences^4^. Cryo-EM has visualized stable D-loop intermediates at equilibrium^29,30^. However, fundamental questions remain unresolved: understanding the structural basis for the 8-nt specificity, dissecting the role of RecA’s CTD in early homology search, and identifying structural determinants of dsDNA capture or rejection.

We address these questions by establishing roadblocked RecA filaments as a system to study homology search, employing time-resolved cryo-EM to capture structural snapshots at 250 ms intervals with varying lengths of sequence homology. Our results indicate that RecA-family recombinases first capture the dsDNA at site II, followed by a transition into three-strand homology sampling by the opening of the dsDNA. We show that the homologous strand retains significant flexibility to compete for base pairing in the three-strand intermediate structure, which is overcome by more than 8-nt of homology. We suggest that the complementary strand is stably bound by site II after homology is fully bound, presenting a hallmark for completed homology search.

## Results

### Data-driven strategy for optimising time-resolved sample preparation

The quality and repeatability of sample preparation is critical for cryo-EM due to the high cost and time required to collect and process datasets^31^. We designed and implemented a sample preparation device as well as a new workflow for rapid data-driven optimisation of cryo-EM sample preparation (Fig 1).

**Figure 1.**
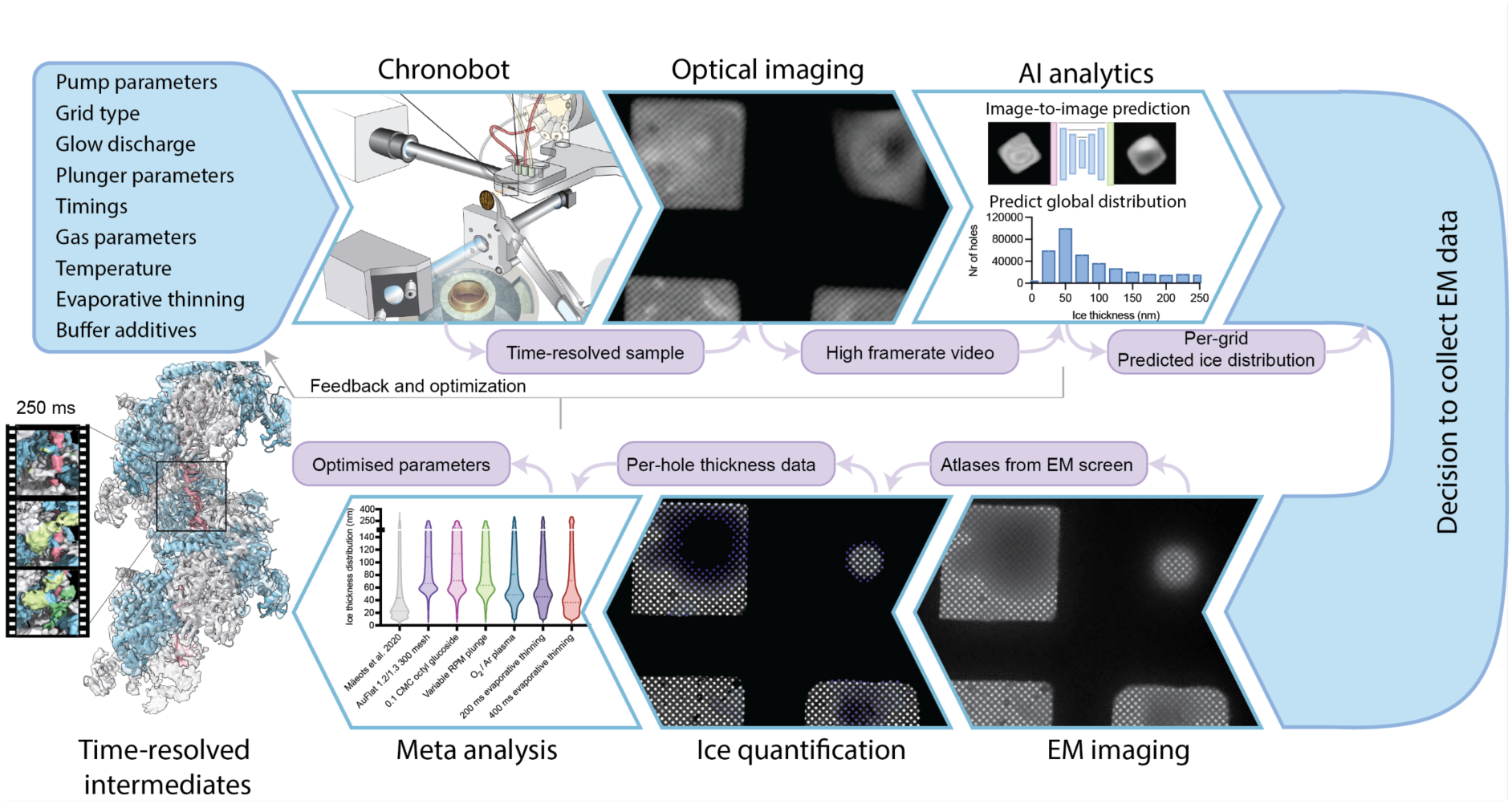
Chronobot workflow overview. From top left: The Chronobot device has several input parameters which can be set per experiment. Each grid is imaged by a high-speed camera before vitrification. Video frames are analysed by a deep learning algorithm to assess overall sample quality. This analysis not only enables an informed decision of whether to collect EM data from the grid but also feeds back data to improve initial sample preparation parameters. If EM data is collected, analytical algorithms are used to estimate ice thickness from EPU atlases. The per-hole thickness data is used for quantitative comparisons between grid conditions over time. This analysis is used for more fine-grained optimisations of conditions. This optimisation cycle is used to rapidly improve grid freezing protocols and can be leveraged to make bespoke protocols for each biological sample that ensure many collectable areas in TEM and, ultimately high-resolution structures.

There are a combination of improvements. The first is a new device focused on robust operation. This includes mechanical improvements to allow finetuning of input parameters beyond what was previously possible. This is supported by electronic feedback and sensor systems to ensure experimental logging as well as monitoring systems that alert users if the Chronobot is not operating at the intended settings.

The second improvement is a combination of two ice thickness quantification algorithms. Both operate at grid-level, providing estimates for the ice thickness distribution as well as quantifying the amount of grid area that is covered by a sufficiently thin vitreous (‘ice’) film. The first operates concurrently with grid freezing and videos are collected automatically each time a grid/sample is plunge frozen. The videos are converted to ice thickness estimates by a machine vision model, providing a coarse estimate of grid quality in real time. This is advantageous in the initial screening process for a new sample or methodological development, allowing rapid elimination of grids with few viable areas or suboptimal ice, and saving days of iterative freezing and screening.

The second algorithm runs concurrently with the standard screening workflow, e.g. atlas collection in EPU (ThermoFischer Scientific), providing a more accurate ice-thickness measurement without requiring extra user time. This enables fine-tuning of parameters to further optimize sample preparation. This workflow allows us to support a wide variety of biological applications of the Chronobot device, providing bespoke solutions to samples where required. In the following sections we outline the details of each of these improvements. Finally, we present one application of the Chronobot by capturing RecA homology search intermediates with various substrates at a 250 ms timepoint.

### Layout of an integrated microfluidic chip and nozzle

The Chronobot device (Fig 2a) is the next-generation improvement of the one described in our earlier work^3^. It remains centred around a microfluidic chip, which has been redesigned to integrate the tasks of mixing, incubating, and then spraying any combination of two reactants (Fig 2b). This modular design functions as a biochemistry-on-a-chip incubator, integrated seamlessly with fluidics, sample handling, and precise mechanical and software control systems. The mixer uses a static 3D serpentine structure to repeatedly break up and merge the channel containing both reactants, which ensures they are mixed within ∼3 ms. For details on the design and validation of the mixer refer to prior work^3^. This is followed by a delay line, where the mixed reactants travel at a fixed speed until reaching the nozzle, allowing for precisely controlled incubation times for the reaction^32^.

**Figure 2.**
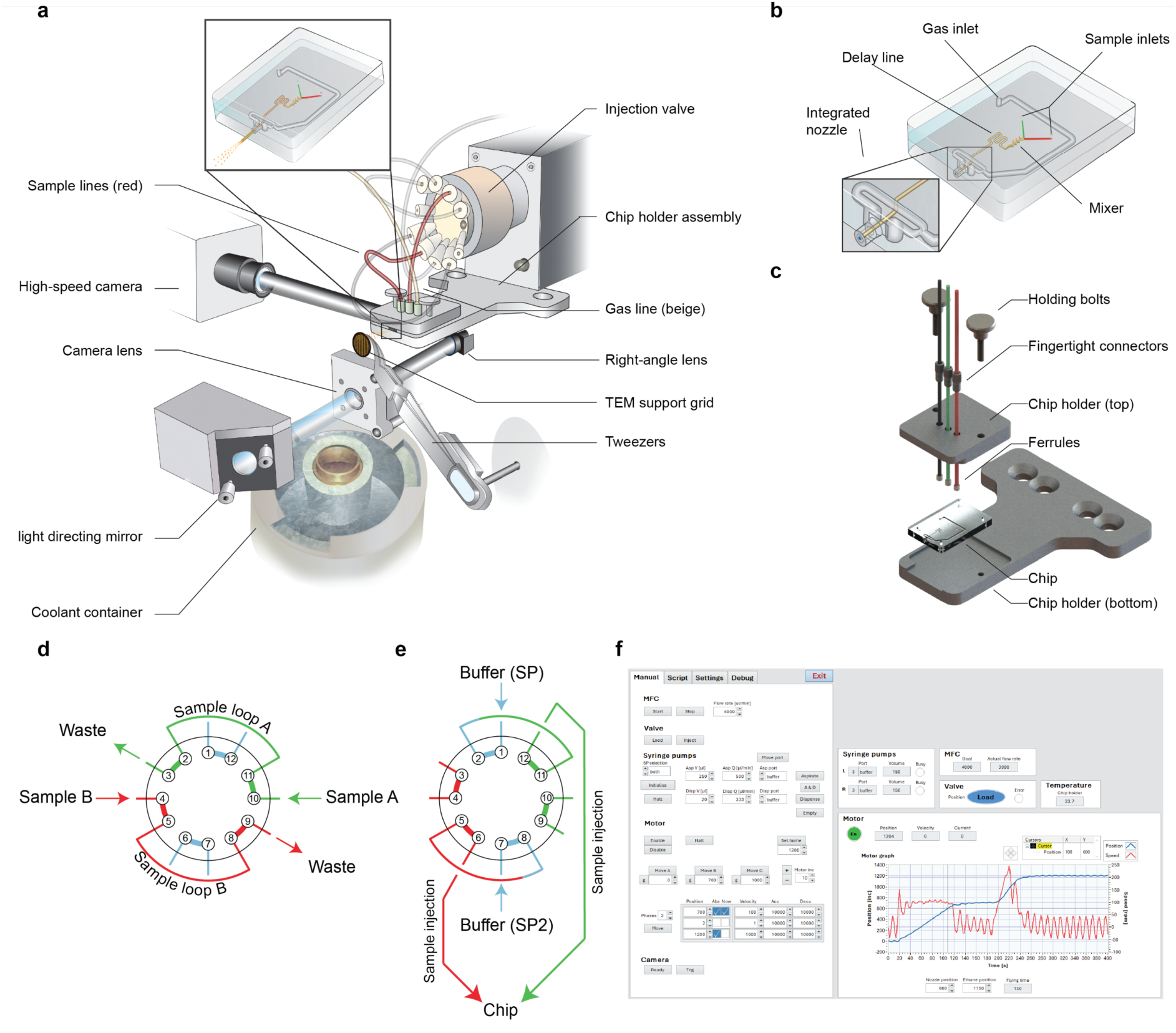
Construction and design of the Chronobot device. **a**, Schematic of Chronobot design. Components are labelled and their relative positions are illustrated. A photo of the operational system is shown in Extended Fig. 1a. **b**, Schematic of the microfluidic device. Samples enter the chip and are passively mixed by being pumped through a mixer structure. This is followed by a variable length delay line that determines the time point. The gas line joins the mixed sample in an integrated nozzle structure, which breaks up the sample stream into small, accelerated droplets. **c**, Chip holder assembly. The holder aligns the chip with the trajectory of the TEM grid, and the inlet tubing with the chip’s ports. **d**, Loading position of the injection valve. Ports 4 and 10 are connected to sample loops for manually loading the sample. **e**, Inject position of the injection valve. Buffer flow from the syringe pumps (SP) pushes sample into the microfluidic chip and initiates the reaction. **f**, User interface for manual control of the system.

The original design relied on PDMS-based microfluidic devices that allowed for rapid and cost-effective prototyping. However, that approach had important limitations. PDMS chip manufacture is labour-intensive and prone to variability in device quality in terms of alignment of the features, thickness of the PDMS layers, and strength of the bond between layers. Another difficulty of using PDMS was that its surface adsorbs proteins, leading to loss of apparent concentration on the grid and uncertainty in the stoichiometry of the reaction. Lastly, we used each PDMS chip a single time, as they cannot withstand the harsh chemicals required for thorough cleaning between runs. This limitation introduced variability in nozzle spraying performance because each new chip required manually cutting borosilicate tubing to fit into prefabricated plastic nozzles, ultimately necessitating multiple replicates per experiment^3^.

To address the above shortcomings, we designed a novel microfluidic device formed out of two distinct layers of etched glass (Fig 2b) produced using the femtoprint method^33^. The nozzle is entirely contained within one layer, while the mixer and incubation lines are split across two layers. The integrated nozzle enables a concentric gas stream with a diameter of 0.9 mm focused around the exit tunnel of the incubated sample which has a diameter of 0.1 mm (Extended Fig 1b). As the sample exits the chip it is broken up into a conical spray with variable droplet diameter and accelerated to high speed which promotes the droplets landing as flat as possible on the cryo-EM support grid. The microfluidic device further features circular cross-section incubation lines, enabling more uniform flow velocity profile compared to the rectangular channels of the PDMS chips, reducing the detrimental effects of laminar flow on time-resolution discussed previously ^2^. The device is mounted in a chip holder which ensures that the tubing is centred around the inlets and can be tightened using fingertight connectors (Fig 2c).

### Sample injection and plunge-freezing

In the previously reported system^3^, samples were directly loaded into the respective syringe pumps, leading to an unnecessarily high sample consumption because the entire 30 µl volume of the system had to be filled with sample. In the Chronobot design, protein samples never enter the pumps during an experiment. The two pumps are instead connected to an injection valve (Fig 2d-e). The injection valve allows the user to inject their protein samples using a syringe into a sample loop, which is attached to the valve. The injection valve has two positions: load and inject (Fig 2d-e). In the load position, the pumps are connected directly to the microfluidic device, bypassing the sample loops containing the protein (Fig 2d). In the inject position, the pumps push buffer through the sample loops and into the microfluidic device (Fig 2e). This injection method allows less sample to be used overall (2 µl per channel per grid), as the system does not need to be filled with the protein samples to function. Furthermore, by placing the injection valve immediately before the microfluidic device and using smaller diameter tubing, dilution and sample loss are reduced, doubling the apparent protein concentration over the previous design.

Gas delivery is now controlled by a mass flow controller valve, fed by a main gas supply. Compared to an externally controlled gas supply, electronic control and sensors are now close to the microfluidic device to ensure a consistent gas flow is maintained during the experiment. The first iteration of the device used a servo motor to control sample collection and freezing.

Despite being an easy to implement and cost-effective solution, its acceleration and deceleration were not tuneable, and maximum velocity was only adjustable in a small range. Moreover, the performance tended to degrade over time. This prompted us to implement a brushless direct current (BLDC) motor, allowing precise control of acceleration and deceleration rates at all stages of grid plunging, as well as output sensor readings regarding the position and velocity of the motor at all points in the experiment. This enabled the grid to travel at different velocities when passing the sample spray, the camera lens, and entering the ethane. A series of XYZ stages permits the precise position of components relative to each other for optimal performance (Extended fig 1a). The distance of the grid from the nozzle (8 mm) was chosen such that the diameter of the spray cone at that distance is ∼4 mm, which fully covers the 3 mm cryo-EM support grid while allowing for a small amount of error in aligning the tweezer tip to the nozzle centre.

### Synchronisation and control of the Chronobot

Precise timing and synchronization between system components are essential for reliable time-resolved sample preparation. Initially, we used an Arduino microcontroller, which allowed rapid and cost-effective prototyping of various configurations^3^. However, the limited software support and closed ecosystem significantly restricted our ability to implement complex software features, limiting reproducibility and advanced experimental capabilities.

In the Chronobot system, we addressed these limitations by designing a custom electronics enclosure that relays information between the PC and system components, providing standardized ports for USB, RS-232, and RS-485 communication, unified into a single USB connection to the PC (Extended Fig 2). The centralized control software, developed using LabVIEW, integrates manufacturer-supported drivers for all system components and provides sensor monitoring and manual control through a graphical user interface (Fig 2f). Users perform experiments via predefined scripts specifying component commands and timing intervals, uploaded directly onto the PCB for automated execution. The software features automatic logging of sensor data and executed scripts, along with built-in error-checking for user-defined scripts.

These advancements significantly enhance reproducibility, data documentation, and overall experimental robustness. More importantly, the improved synchronization and precise mechanical control provided by the new electronics and software are foundational to advanced functionalities described later.

### Quantification of cryo-EM sample quality

To optimise sample preparation workflows for time-resolved cryo-EM, we first need to define a metric by which to quantify produced samples. High-quality cryo-EM samples are commonly understood to contain large areas of thin ice, with densely packed but monodisperse protein. In our workflow, we currently aim to quantify and optimise the former. Usually what is meant by thin ice is vitreous areas with high contrast particles visible, which we empirically found to be around 60 nm thickness or below. This thickness also corresponds to the highest resolutions obtainable in the final reconstruction^15^. We therefore adopted the number of holes on a cryo-EM support grid which contain ice below 60 nm, as our major optimisation metric. This metric has the benefit of distilling sample quality to a single comparable value, which aids our workflow. In order to capture additional effects not captured by this metric, we also use ice thickness distributions as a secondary readout.

### Rapid ice thickness estimation using deep learning

The integration of the high-speed camera into the Chronobot workflow enables instant feedback to the user about the general appearance of the sample (Fig 2a and 3a). However, to integrate this imaging into the workflow more efficiently we aimed to quantify per-pixel ice thickness from these images. Capturing videos while plunging imposes significant limitations on the fidelity of the imaging. Since the imaged grid is in motion, the videos exhibit variation in focusing across the grid surface. Imaging transmitted light through the grid limits the brightness of the captured images. Due to these limitations and the relatively low spatial resolution obtainable, we opted not to pursue an analytical solution to ice thickness estimation in this imaging mode.

**Figure 3.**
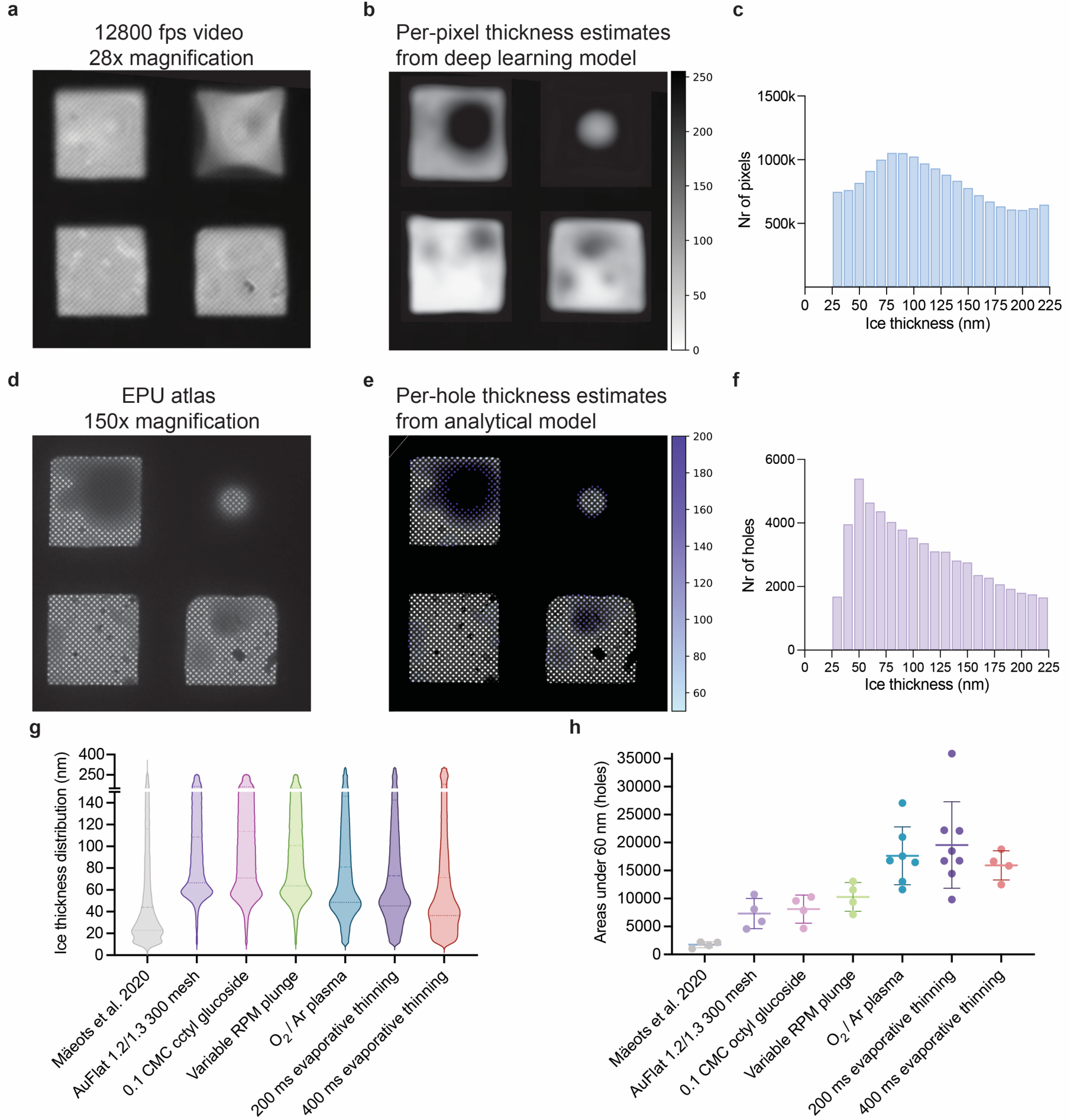
Quantification of ice thickness. **a**, Representative image from high-speed camera video frame. **b**, Example of deep learning model outputs. Since the model works on individual grid squares, they are manually aligned here for visual comparison with **a**. **c**, Example of grid-level prediction of ice thickness distribution, as determined by combining all inferred squares on a grid. **d**, Representative micrograph taken at the 150x magnification on a Talos Arctica, which is used as input into our analytical model. **e**, Example of per-hole predictions determined from d. **f**, Example of grid-level prediction of ice thickness distribution by combining all predicted atlas tiles. **g-h**, Comparison of different grid preparation parameters using ice thickness distributions or our <60 nm heuristic, respectively. All conditions except for varying evaporative thinning times are cumulative with previous improvements.

Image processing by deep learning has been an area of major development in cryo-EM in recent years. This includes applications at all levels of the pipeline, from sample preparation, micrograph denoising, particle picking, to final reconstructions^34–37^. This is principally due to the ability of deep learning models to detect patterns in input images which contain substantial noise. Here we present an in-house model for the estimation of ice thickness from high-speed camera movies (Extended Fig 3g). The model is based on the DeepLabv3 architecture, which is an image segmentation model, with features specifically designed to increase field of view and incorporating multi-scale context^38^. The atrous spatial pyramid pooling (ASPP) approach of DeepLabv3 integrates information from several discrete fields-of-view at different sampling rates^39^, in contrast to standard 2D convolutions use invariant local image transformations. This model was chosen because the correlation between pixel values in the videos and ice thickness appeared to be mostly affected by the shape and size of the sample droplet, rather than local intensity. Therefore, we aimed to create a model which would be more attuned to the context of an entire droplet. We report a custom decoder that replaces the previous DeepLabv3 decoder, to specifically accomplish the ice thickness regression task rather than segmentation. The model was trained on pairs of images comprising individual grid squares. The input images were grid squares segmented from the video frames, while the target images were thickness-encoded images of the same areas (Extended Fig 3j-l). How these images were generated is discussed in detail in the subsequent sections. Training and evaluation metrics are reported in Extended Fig 3h, as well as the performance of the model on per-pixel thickness estimation (Extended Fig 3i).

The model outputs an overlay of the grid squares in real-time, with inference running simultaneously as the user operates the Chronobot (Fig 3b-c). Using this coarse estimate, it is possible to rapidly detect problems with the samples such as support grids being too hydrophobic, suboptimal spray density on the grid, broken support films, or effects of the sample solution. In addition, users can run a set of conditions and rapidly determine the most promising ones without screening the grids in a TEM. For samples which are imaged using TEM, we then perform a fine-grained ice distribution estimate using analytical methods described below.

### Ice thickness estimation from low magnification TEM images using aperture limited scattering

We present a method for quantifying screened grids in terms of ice thickness distribution as well as the total number of collectable areas in an unbiased way (Fig 3d-f). Our approach is an expansion of previous work^18,19^. The approach relies on aperture-limited scattering (ALS), where scattered electrons by the ice are filtered out by the objective aperture of the microscope.

Thicker ice scatters electrons more strongly, which move outside the aperture and don’t contribute to image formation. This effect makes thicker ice appear darker in the final image. The first requirement for this method is to collect a calibration dataset for the microscope, using rigorous ground truth measurements of ice thickness. We chose the ice-channel method for our ground truth data (Extended Fig 3a)^17^. This is then calibrated with 150 000x magnification images, with a reference vacuum intensity image. This allows us to fit a model for the ALS coefficient of our screening microscope:

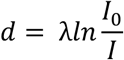

Where *d* is ice thickness, λ is the ALS coefficient, *I*_0_ is the intensity of the image over vacuum, and *I* is the observed image intensity over the ice which is to be measured. In practice using ALS for ice thickness estimation is accurate as long as *I*_0_ can also be measured within the same image, and λ has been accurately estimated by calibration (Extended Fig 3b)^18^.

Previous applications of this method were intended to aid the user by using ice thickness measurements to choose areas for data collection in real-time. Since our goal is to obtain grid-wide distributions of ice thickness, we chose to work at the ‘atlas’ magnification of our microscopes, which is 150x. To achieve this, we collect a dataset of micrographs taken at 150 000x magnification and estimated their ice thickness using the ALS method. We then use these as ground truth for fitting the ALS coefficient at 150x magnification. Although this results in compounding errors from the two-step calibration, we could collect a larger dataset that would not have been obtainable using the more accurate ice-channel method. This larger dataset makes the model fit much more robust to the inherent noise in low magnification images (Extended Fig 3c).

We then applied this ALS model in a data pipeline which takes as input the atlas tiles collected at 150x magnification and estimates hole locations across the entire grid (Extended Fig 4). In addition to hole location estimation, the software also estimates vacuum intensity across each tile by fitting a simple polynomial model on the brightest points seen in each image. This allows for the best estimate of *I_0_* across the image. The algorithm also includes several filtering steps to exclude contaminants, broken foils, and areas of ice too small to be included in a realistic data collection session. The output of the model is a per-hole estimate of ice thickness covering every visible hole on the grid (Fig 3d-f).

**Figure 4.**
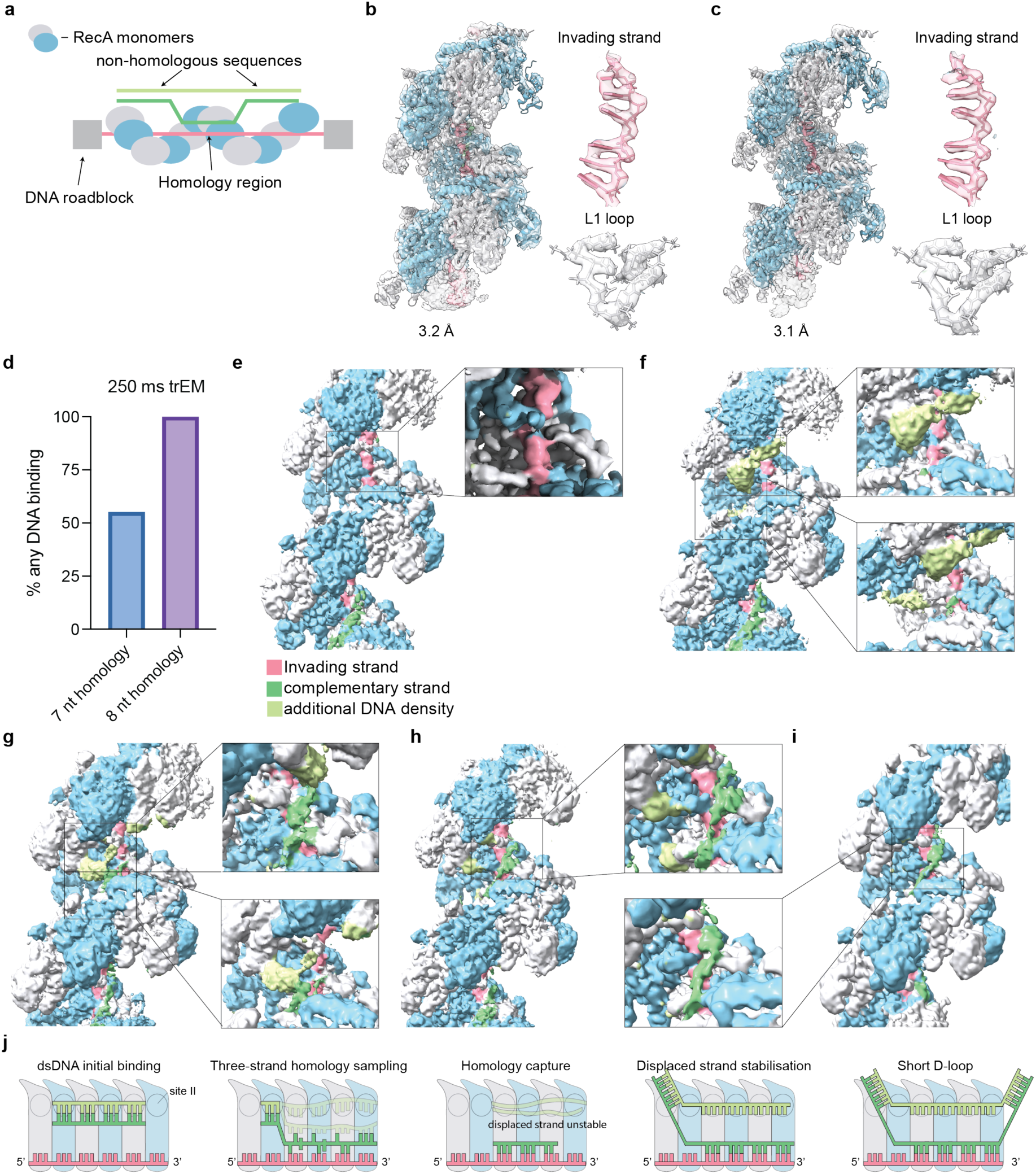
Time-resolved structural studies of RecA homology search at 250 ms. **a**, Schematic depiction of the roadblocked mini-filament structure. **b and c**, Consensus refinements of (RecA^mini-filament^+dsDNA^8-nt^)^250ms^ and (RecA^mini-filament^+dsDNA^7-nt^)^250ms^, respectively. The L1 loop and invading strand densities are depicted to represent the quality of the maps. PDB-7JY9^29^ was docked into both densities. **d**, Quantification of observed DNA binding between 7-nt and 8-nt substrates. DNA binding is defined as any densities observed at the CTD, site II, or complementary strand positions. **e**, Dissociated conformation in (RecA^mini-filament^+dsDNA^7-nt^)^250ms^. All classes display complementary strand density at the 3’ end of the mini-filament, due to dsDNA under the HpaII roadblocks. The RecA filament partially forms on this stretch of dsDNA before the roadblock. **f**, dsDNA capture conformation in (RecA^mini-filament^+dsDNA^8-nt^)^250ms^. **g-h**, Putative three-strand intermediate conformations in (RecA^mini-filament^+dsDNA^8-nt^)^250ms^. **i**, Homology bound conformation in (RecA^mini-filament^+dsDNA^8-nt^)^250ms^. **j**, Proposed model for sequence of events during RecA-mediated homology search.

### Systematic improvements of ice quality for time-resolved cryo-EM

We apply our rapid ice thickness estimation methods to systematically improve every aspect of time-resolved sample preparation (Fig 1 and Fig 3g-h). As a baseline we apply the protocol reported in our previous work (Fig 3g-h, ‘Mäeots et al. 2020’)^3^. The first major improvement we implement is using gold grids^40^. Using 1.2/1.3, 300 mesh AuFlat grids (Protochips) results in a 318 % uplift in the number of collectable areas (Fig 3h ‘AuFlat 1.2/1.3 300 mesh’). We observe a significantly flatter ice distribution on gold foil compared to carbon foil (Fig 3g ‘AuFlat 1.2/1.3 300 mesh’). While average ice thickness appears higher over gold, thin areas on carbon foils are extremely rare and thus up to 6 grids were needed to collect even a modestly sized dataset previously^3^(Fig 3h). Therefore, we adopt AuFlat 1.2/1.3 grids for all subsequently tested conditions. The second parameter we optimise is the addition of a detergent, octyl glucoside (OG). Using this detergent at 1/10 critical micelle concentration (0.1 CMC), we observe an 11 % increase in collectable areas with a more uniform thickness distribution (Fig 3g-h ‘0.1% Octyl glucoside’). OG detergent at 0.1 CMC was adopted for all conditions described below.

Thanks to the introduction of a BLDC motor, we can vary the speed of the grid at multiple points during the plunging motion. We find moving past our droplet spray at 80 rpm to enable an optimal balance between grid coverage and foil damage. After collecting the droplets, we slow the motor down momentarily in front of the high-speed camera, followed by a rapid reacceleration to ensure efficient vitrification. An example of the plunging profile is shown in Extended fig 1e. We observe a 27 % increase in collectable areas by using variable rpm plunging over constant rpm plunging (Fig 3h ‘Variable RPM plunge’).

Unlike blotting methods, sample preparation by spraying is highly dependent on the hydrophilicity of the support grid as a thinning mechanism. We experiment with different glow discharge and plasma cleaning conditions to improve the ice distribution. We found that oxygen-argon plasma at 40 W for 1 min was ideal for achieving thin ice, improving the number of collectable areas by a further 72 % (Fig 3h ‘O_2_/Ar plasma’).

Finally, we test evaporative thinning as a potential method for achieving even higher quality ice. ^41^This involves tuning in additional time between collecting the droplets arriving on the grid and being plunge-frozen. During this time the liquid layer on the support grid thins rapidly. Although this requires additional time in the time-resolved experiment, we found that this method can achieve the thinnest ice and achieve an additional 11% improvement in collectable areas and can be compensated using shorter delays in the microfluidic device (Fig 3g-h ‘200 ms evaporative thinning’).

By combining all reported improvements (Fig 3g,h), we are now able to produce cryo-EM samples that have on average ∼20k holes covered by ice which provides excellent contrast and does not limit achievable resolution^15^. This represents an average 11-fold improvement over the initially reported protocol^3^. Below we apply the optimised Chronobot method to study homology search in RecA.

### Assembly of roadblocked RecA filaments

We next applied the Chronobot workflow to the initiation of RecA-mediated homology search, a fundamental aspect of HR^25^. Single-molecule studies^4^ revealed a crucial kinetic difference during dsDNA homology recognition: RecA stably captures dsDNA containing at least 8 nucleotides (nt) of microhomology with residence times on the order of seconds, whereas shorter sequences (≤7 nt) bind transiently, with residence times of milliseconds. We hypothesized that trEM and single particle analysis could directly capture and visualize these transient molecular interactions, thereby clarifying the mechanism of early homology search and recombinase sequence specificity.

To further facilitate this study, we design and generate short, roadblocked RecA filaments amenable to single-particle cryo-EM analysis. RecA filaments were capped at both ends by covalently attaching methyltransferase HpaII proteins as steric roadblocks on the ssDNA, preventing filament elongation (Fig 4a, Extended Fig 5a-b). This design maintains a native-like filament structure while also spatially restricting homology search interactions precisely to the filament’s midpoint (Extended Fig 5c), making it amenable to particle alignment in single particle analysis. The resulting roadblocked mini-filaments demonstrate structural uniformity and integrity (Extended Fig 5d-e). The filaments consist of 12 (with a small subpopulation of 13) wild-type RecA monomers assembled around single-stranded DNA in the presence of ATPγS (Extended Fig 5f).

### Three-strand dynamics govern RecA homology search at millisecond timescales

We collect time-resolved data using two different dsDNA binders, containing a single 7- or 8-nt binding site flanked by non-homologous sequence (Extended Fig 5c). The RecA mini-filaments are mixed with 3-fold excess of the chosen dsDNA inside the microfluidic device. An excess of dsDNA was chosen to ensure that interactions with the RecA filament were not constrained by DNA concentration at 250 ms. The two datasets are hereafter referred to as (RecA^mini-filament^+dsDNA^8-nt^)^250ms^ and (RecA^mini-filament^+dsDNA^7-nt^)^250ms^. Consensus reconstructions from both substrates yield high-quality single particle cryo-EM reconstructions (3.2 and 3.1 Å resolution respectively) resembling equilibrium RecA structures (Fig 4b-c)^29^. At 250 ms, the 7-nt substrate exhibited ∼50 % occupancy, consistent with its reported half-life (Fig 4d-e)^4^. In contrast, the 8-nt substrate shows nearly 100% occupancy, reflecting its longer residence time (Fig 2d)^4^.

Local 3D classification of (RecA^mini-filament^+dsDNA^8-nt^)^250ms^ and (RecA^mini-filament^+dsDNA^7-nt^)^250ms^ reveals multiple distinct configurations (Extended Fig 6, Extended Fig 7). An initial dsDNA capture state features unopened dsDNA engaged at RecA’s secondary DNA binding site (site II or S2), a previously hypothesized but unobserved structural state under equilibrium conditions^41–43^ (Fig 4f). This state displays no density for the complementary strand bound at site I but has a strong density at site II for two consecutive monomers (Fig 4f, top inset). This is accompanied by a novel DNA interaction between site II and the RecA C-terminal domain (RecA^CTD^), where dsDNA does not contact site II of the same monomer that binds the RecA^CTD^ (Fig 4f, bottom inset). This is distinct from interactions reported in D-loop conformations obtained at equilibrium using stabilising DNA constructs that show dsDNA binding to RecA^CTD^ only. Due to the limited local resolution of this density, we are unable to assign with certainty whether it corresponds to dsDNA or ssDNA, however the contact with L2 loop bares similarity to the reported dsDNA structures^29^.

Additional intermediates are consistent with progressive homology sampling (Fig 4g-h, Extended Fig 8c). Intriguingly, despite clearly resolving L2 loops, we do not observe continuous displaced strand density at site II, as seen in late equilibrium states^29,30^. We report an apparent three-strand intermediate, characterised by a strong density perpendicular to the invading strand making contact between site II and the L2 loop, along with density for the complementary strand towards the 3’-end of invading strand (Fig 4g, top inset), and an additional density between two adjacent L2 loops in the 5’ direction (Fig 4g, bottom inset). This additional density is not making a clear contact with the invading strand, indicating it does not belong to the complementary strand. Additionally, we observe an intermediate where the displaced strand density is only present proximal to the L2 loops and not along site II (Fig 4h, inset). This observation supports previously suggested transient three-stranded intermediates^27^ and suggests that stable capture of the displaced strand at site II could be a hallmark of a completed homology search. We do not observe ordered triplets at the complementary strand position in this conformation, supporting this notion (Fig 4h, inset).

We also observe a homology bound state of (RecA^mini-filament^+dsDNA^8-nt^)^250ms^ (Fig 4i). This conformation displays ordered triplets bound at the predicted homology location (Fig 4i, inset). However, we do not observe stable capture of the dsDNA ends at RecA^CTD^ along with this conformation, nor do we observe the stable capture of the displaced strand at site II. This indicates that either stable homology binding precedes both events, or that the 8-nt homology dsDNA construct used here is not long enough to stabilise these binding events.

We investigate the dynamics of CTD binding within our datasets (Extended Fig 8 a-b). Previous cryo-EM studies of homology search suggested that binding of dsDNA to sequential CTDs of RecA was a probabilistic mechanism of homology search, with the likelihood of dsDNA unbinding increasing with each step along the filament^29^. In our dataset, the likelihood of dsDNA being bound to any RecA^CTD^ in the filament is 31% for (RecA^mini-filament^+dsDNA^7-nt^)^250ms^ and 34 % for (RecA^mini-filament^+dsDNA^8-nt^)^250ms^ (Extended Fig 8a). Several potential dsDNA binding modes to site II are also identified (Extended Fig 8c).

To follow the observation that overall DNA occupancy is much lower in the 7-nt dataset, we also observe the structure and stability of the homology binding. By comparing the most stable binding states in both conditions, we observe a significant increase in structural order of the complementary strand in the 8-nt conditions (Extended Fig 8d, left). In this case, multiple classes resolve two stable triplets of homologous DNA, albeit at lower local resolution compared to the invading strand (Extended Fig 8d, left). In contrast, (RecA^mini-filament^+dsDNA^7-nt^)^250ms^ does not contain any classes where individual DNA bases could be resolved in the complementary strand, despite similar global resolutions between classes (Extended Fig 8d, lower left vs right). This suggests that the backbone and base pairing in this condition are not stable, consistent with the previous observation that the apparent oU-rate of this complex is significantly higher than the 8-nt homology complex^4^.

## Discussion

### Improved sample preparation device

This work introduces the Chronobot, a device for preparing time-resolved cryo-EM samples, as well as a new workflow for optimising sample preparation in cryo-EM by leveraging machine learning at multiple levels of grid-level ice thickness quantifications. A key feature of the Chronobot system is the modular nature of its microfluidics. Devices with various delay lines can be easily replaced into a user-friendly chip holder permitting access to multiple time-points in the millisecond to second range. These new devices are manufactured from glass and thus are readily washable and reusable, significantly cutting down on manual labour required to manufacture PDMS alternatives^3^. The integration of the nozzle structure into the glass device provides a predictable spray cone with high uniformity. This contrasts with previous spraying methods which introduced significant variability across replicates^3^. Future work in this area will introduce further functionality to the device such as multiplexed delivery of different timepoints of a reaction simultaneously onto the support grid.

The introduction of a sample injection valve reduces the minimum system volume needed to operate the system from 6 μl to 2 μl. Future work will incorporate spectrometry measurements to precisely quantify the elution of the sample from the microfluidic device, which would enable reducing sample consumption further and reach peak concentration on the grid.

### A quantitative approach to sample preparation optimisation

The Chronobot approach is designed to rapidly optimise grid preparation protocols by quantifying sample quality and thus enabling a reproducible and reliable sample preparation workflow. This enables a wide variety of biological samples to be studied, rapidly and inexpensively optimising conditions for each sample with high confidence. As an example application, we show how adjusting the speed of the motor at several steps during the plunging motion can improve final sample quality. We also show how individual parameters such as plasma treatment of grids prior to sample freezing can be successfully optimised using our workflow. With the introduction of evaporative thinning, ice thickness can now be finetuned into the system, albeit with the potential limitation that additional reaction time is added Notably, during the final stages of preparation of this manuscript, an independent study with different experimental setup and conditions reported a conceptually similar evaporative thinning approach with equally positive outcomes^44^.

By combining various optimisations across several areas: mechanical design, electrical design, sensors, microfluidic devices, sample delivery and software control we can reliably produce time-resolved samples with, on average, ∼20k high quality collection holes per grid. In our empirical observations, after having optimised the Chronobot settings for a given sample, a single vitrified grid provides sufficient areas for collection of a high-resolution time-resolved dataset, thus obviating the need for many technical replicates. This improvement enables more time points and reaction conditions to be prepared using the same amount of sample and experiment time as before. Future work will be aimed to make the Chronobot device and workflow more readily available to the community.

### Visualising time-resolved RecA homology search initiation intermediates

RecA-mediated homology search is underpinned by rapid interactions with dsDNA substrates. The latter are bound, opened, and scanned by a presynaptic RecA filament within a few hundred milliseconds, resulting in either a dissolution of the complex or stably bound states^4,25,29^. Since the initiation steps of this process are out of reach for standard cryo-EM sample preparation technology, we reasoned that it would represent both an important topic of major biological relevance as well as more broadly validate the capabilities of the Chronobot system. Here we report the direct visualisation of reaction intermediates with half lives in the millisecond regime using the Chronobot system.

Recent studies of RecA- and its human homologue Rad51-mediated strand-exchange have uncovered the structure of the D-loop at equilibrium using stabilising substrates^29,30^. These structures illuminated the dsDNA-binding functions of the RecA^CTD^ and showed how that dsDNA can be captured by different monomers in a dynamic equilibrium. Meanwhile the homologous and complementary strands were seen stably bound to the invading ssDNA and site II, respectively^29^. Despite these advances, it remained unclear how the dsDNA is captured and opened during the initial, pre-equilibrium, stages of this process. It was further unclear how this homology search would differ when less than the required length of homology is present, as it is known to not form a stable D-loop structure. Using trEM, we present several intermediates of homology search at 250 ms that precede D-loop formation and complement previous studies.

To our knowledge, this is the first report of a homology search intermediate that depicts dsDNA capture without homology binding or local opening of dsDNA, thus visualising the initiation of homology search. Surprisingly, dsDNA binding appears to occur at site II, which has previously been postulated^41–43^ but not observed (Fig 4j, “dsDNA initial binding”). At this early time point, RecA^CTD^ binding of dsDNA was not observed in the absence of homology binding, which we suggest could further support a role for the RecA^CTD^ in strand-exchange. We also report several related conformations to the site II dsDNA capture state (Extended Fig 8c). Based on this we suggest a transformation of the dsDNA from site II into a three-strand intermediate conformation (Fig 4j, “Three-strand homology sampling”). We observe that short homology can be fully bound without clear density for the displaced strand at site II or the RecA^CTD^, which we propose might represent the end of three-strand sampling where the homologous strand is out-competed by the invading strand but not stabilised (Fig 4j, “Homology capture”). Based on the data presented, we propose that stable capture of the displaced strand at site II could be a hallmark of successful homology capture, as we do not observe continuous density at this site in our experimental conditions (Fig 4j, “Displaced strand stabilisation”). This would include the formation of the short D-loop structures reported previously^29^, which we do not observe in our experimental condition (Fig 4j, “short D-loop”). This model of homology search initiation is likely more linear than the true process since we do not account for all the observations presented. A key open question remains with dsDNA binding to the RecA^CTD^ which we observe in several classes but were unable to assign a function with available data. Future work will be required to fully delineate the functionally relevant states by studying additional lengths of homology at more time points of the reaction.

We also address the differences between 7-nt and 8-nt of microhomology, which has previously been reported to be a key feature of RecA-family recombinases, where dsDNA substrates featuring continuous homologous sequences of 7-nt or fewer are rapidly rejected while longer ones remain stably bound^4^. At 250 ms, we observe stark differences in the structural order of the bound homology between our two dsDNA substrates. We observe that in the 8-nt case, all eight bases can be clearly defined in some classes, including the canonical stretched backbone between adjacent triplets (Extended Fig 8d, left). In contrast, such ordered density was not observed with the 7-nt substrate, indicating that base pairing and the DNA backbone are not stable in this case (Extended Fig 8d, right). This is further supported by our quantifications, which show that around half of RecA mini-filaments are in a dissociated state at 250 ms using the 7-nt substrate, while all mini-filaments are DNA bound when mixed with an 8-nt substrate (Fig 4d). These observations support previous findings regarding homology binding kinetics^4^.

In our time-resolved datasets, we do not observe the large variability in RecA^CTD^ capturing dsDNA reported previously at equilibrium^29^. Further work could explore whether such a population of binding states would develop during longer time points and in the presence of a longer dsDNA substrate that would be competent for complete strand exchange.

A broader open question relates to how much of this mechanism is conserved in other RecA-family recombinases like Rad51^30^. Although previous data suggest that the sequence preference for homology of at least 8-nt in length is shared between Rad51, Dmc1, and RecA^4^, it would be enlightening to observe how the initial Rad51 homology search dynamics change due to the lack of an equivalent C-terminal domain. Other major questions relate to various modulators of Rad51 activity and how this process proceeds in a chromatin context^45^.

As exemplified by the single particle analysis of time-resolved RecA datasets reported here, comprehensive structural analysis of time-resolved datasets remains an open challenge. They likely contain more heterogeneity and potentially resolvable states than can be effectively extracted with currently available methods, motivating continued work on the efficient processing of such data^46^. In summary, the time-resolved method presented in this work is generally applicable to biochemical processes occurring on millisecond timescales. Thanks to advancing technological development^2,3,47–57^, time-resolved cryo-EM will increasingly and more commonly allow deep mechanistic understanding of many processes as well as aid the early stages of structure-based drug discovery.

## Materials and Methods

### Deep learning methods for ice thickness estimation

#### High-speed camera imaging

High-speed camera images were acquired with Nova S12 type 1000 K (Photron) and a LED light source (F5100, Photonic). Lenses used were a 2x fixed lens, adjustable 0.58x-7x lens, and a front 2x attachment lens (Navitar). Camera control was achieved by the manufacturer’s software (PFV4, Photron). LED light was directly behind the imaged area for optimal illumination. Standard imaging settings for sprays were 12,800 frames per second, 600 000/s shutter speed, 1024 x 1024-pixel resolution. An optical trigger attached to the tweezer holder synchronises the camera imaging with the motor.

#### Deep learning Dataset

Video files are processed in Python. With visible light imaging, there is always high contrast between grid bars and grid squares, therefore binarisation at a fixed threshold is sufficient to detect locations of grid squares in each image. Array slices are extracted around the centres of the boxes to generate model inputs. EM atlases are aligned to video files by selecting an alignment point in both images and then using several rounds of rotation, zoom, and translation to determine the optimal alignment. Grid squares detected in the visible light image are used to extract the box at the same location in the EM image. In total 45542 grid square pairs were extracted from 20 cryo-EM grids. For training stability, grid squares that had dramatically different appearance in the two datasets were discarded. Some examples of discarded pairs are foil breakage after high-speed camera imaging, strong contamination in the EM image, or very poor focus in the light image. For training and validation, a 95/5 split was used. Augmentations of the dataset included flipping, rotations, and resizing in the range 0.95-1.05 on both axes.

#### U-Net-style encoder–decoder with ASPP

Our ice-thickness network follows an encoder–ASPP–decoder paradigm. It combines a pretrained ResNet-101 encoder for low level feature extraction with an atrous spatial pyramid head for large receptive field context and a decoder that fuses multi-scale skip features while progressively restoring spatial resolution. We initialised the ResNet-101 encoder with ImageNet-1K weights and replaced the first 7×7 convolution with a single-channel variant to accommodate our inputs. Skip features are after layer 1 (C=256, 32×32), layer 2 (C=512, 16×16) and layer 3 (C=1024, 8×8), providing progressively coarser but semantically richer representations. The output of layer 4 (C=2048, 4×4) feeds the ASPP head. The ASPP module uses dilations of 6, 12, and 18 for the atrous convolutions. Branch outputs are concatenated and projected with a 1×1 convolution, batch-normalisation, ReLU and 0.5 dropout, yielding a 256-channel, 4×4 context tensor. The ASPP head supplies global geometric context that is crucial for interpreting whole-droplet morphology, while the skip-connected decoder preserves fine grid-bar boundaries needed for accurate thickness delineation. The second part of the decoder consists of sequential bilinear upsampling and 2D convolution steps, avoiding checkerboard artefacts sometimes observed with transposed convolutions^58^.

#### U-net model training

The hybrid loss function employed here combines two complementary components: a Jensen-Shannon Divergence (JSD) loss and a weighted Mean Squared Error (MSE) loss. The JSD component calculates divergence between smoothed marginal probability density functions of predictions and ground truths, applying Gaussian kernel smoothing ^59^. Due to large parts of the images being either black (gridbars, thick ice) or white (vacuum), loss values are weighted towards middle values by lower scaling on either end to deprioritize these regions. The weighted MSE has the same weighting to prioritize grey values corresponding to ice. Together, these terms encourage predictions that are both distributionally correct and locally precise. For training, the Adam optimizer was used with the OneCycleLR scheduler and training lasted 50 epochs. The model checkpoint with the best validation loss was chosen for inference.

### Analytical methods for ice thickness estimation

#### Automatic estimation of intensity over vacuum (I_0_)

Two factors contribute to the observed image intensity over vacuum. The first is inherent noise in low-dose electron microscopy. The second is non-uniformity in beam intensity across the field of view. The noise in the image was smoothed by applying a circular 2D-convolutional filter. A radius of 5 pixels was chosen, which averages over an area of 0.92 μm diameter at our magnification, which is smaller than the hole size of commonly used cryo-EM support grids. To account for variations in beam intensity, all areas within 95 % maximal intensity are then considered vacuum. This is used to fit a 2^nd^-degree polynomial function across the image the estimate variations in beam intensity. Due to edge effects of polynomial functions hard clipping is employed to limit this background correction to the range −10 to 10 in predicted nm. In practice this provides a faster and superior estimate of intensity over vacuum compared to manual selection of suitable areas.

#### Automatic detection of hole sizes and patterns

Due to lensing distortion of images at low magnification, it is necessary to find holes in each grid square separately rather than fitting a global repeating pattern. This is done by first binarizing the image with a low threshold so that only grid bars are masked out. Then the square outlines of the grid squares are estimated. The size, spacing, and angle of the holes on the support foil are then estimated in a two-step process. First, we generate a mask to find holes, masking out every pixel that corresponds to I0/I > 6, indicating very dark areas. Parameters are then estimated for 100 randomly selected binary areas (holes), measuring their diameter, distance to the nearest other binary area, and the angles between their centres of mass. These initial estimates are then used to fit a hole pattern that covers the entire square outline, searching initially over ± 2 standard deviations of the initial estimates. We found that the most robust method for fitting square patterns involves the generation of a binary image with a black background and white dots of the estimate hole size. These masks are then iteratively sampled at different rotations and offsets to find a binary pattern that best fits the TEM image. For gold foils, this is highly effective due to the dark appearance of the foil in TEM. This process is done iteratively for all grid squares, while progressively narrowing the search ranges as the results converge. All functions are GPU accelerated using CUDA.

#### Ice quantifications

Using a collected atlas of the cryo-EM grid, ice thickness was estimated on each background-corrected atlas tile to gather data on every area of the grid. A thickness map is then created by multiplying the image by the ALS function for ice thickness per-pixel. The ALS coefficient at 150x used is 133.9. To estimate per-hole thickness we employed the 5-pixel circular kernel with a 0.92 μm diameter, however unlike in the vacuum intensity estimation, here a 5% percentile filter is used instead, which replaces each pixel value with the 5^th^ percentile value within the filter radius. This is done as a passive filter against ice contamination. While the gradient of ice thickness across holes is not large in at 1.2 μm, a contaminant partially overlapping the hole can skew measures such as the mean or median. Therefore, we found that looking at the brightest pixels in the image is robust both to noise and contamination. There is a small underestimation of ice thickness that can result from this, which we estimate this to be around 1-2 nm in typical use. This was a key optimisation for obtaining global ice thickness distributions per grid, as sprayed grids can have ∼50% of the total surface area not covered by ice, which makes the total statistics sensitive to any measurement noise over empty areas. The second optimisation we employ to ensure robustness is the clustering of holes using the DBSCAN algorithm. We cluster holes that are deemed to have ice, and then remove any clusters which contain fewer than 8 adjacent holes. Empirically this removes most contamination that has remained after the percentile filtering, as genuine droplets on the grid tend to cover larger areas.

### Preparation of roadblocked RecA filaments

#### Expression of the HpaII methyltransferase

Frozen T7 express competent high-efficiency E. coli cells (NEB) for HpaII overexpression were a kind gift from Hasan Yardimci laboratory. T7 cells are deficient for restriction enzymes that cut methylated DNA. Plasmids containing HpaII were received from the Hasan Yardmici laboratory and modified to include a Twin-strep-tag. 8L of cultures in Luria Broth (LB) medium (1% (w/v) bacto tryptone, 0.5 % (w/v) yeast extract, 0.01% (w/v) NaCl), supplemented with Ampicillin (100 µg/ml) were grown at 37 °C until OD600=0.6 was reached. Protein expression was then induced with 0.4 mM Isopropyl ß-D-1-thiogalactopyranoside (IPTG). Cultures were harvested 2 hours after induction by centrifugation at 4500 rpm for 15 mins at 4 °C and cell pellets snap-frozen in liquid nitrogen.

#### Purification of the HpaII methyltransferase

50 mL of lysis and binding buffer (20 mM Tris-HCl pH 8.5, 500mM KCl, 10mM Imidazole, 10 % (v/v) Glycerol, 1mM tris(2-carboxyethyl)phosphine (TCEP), 0.7 mM phenylmethylsulfonyl fluoride (PMSF), 1x Protease Inhibitor Cocktail, Roche) was added to cell pellets. Cells were lysed by sonication (2 x 2 min on time total; 2 sec pulse; 5 sec oU; Amplitude 40%). Lysate was clarified by centrifugation at 22,000 rpm for 20 min at 4 °C. Cell lysate was applied to a 5 ml Strep-Tactin Superflow plus Cartridge (Qiagen). Column was washed with 50 mL of washing buffer (25 mM Tris 7.8, 250 mM KCl, 5 % glycerol, 1 mM TCEP + EDTA-free protease inhibitor tablets) and eluted with 4 mL elution buffer (25 mM Tris 7.8, 250 mM KCl, 5 % glycerol, 1 mM TCEP + EDTA-free protease inhibitor tablets, 2.5 mM Desthiobiotin). Eluted protein was loaded onto 120 ml HiLoad Superdex 75 16/600 sizing column (Cytiva) and run in gel filtration buffer (50 mM Tris pH 7.8, 150 mM KCl, 1 % glycerol, 1 mM TCEP). Pooled fractions were concentrated to ∼10 mg/ml in storage buffer (50 mM Tris.HCl pH 8.5, 150 mM KCl, 1% glycerol, 1mM TCEP) and frozen at −80°C.

#### DNA labelling with HpaII methyltransferase

Oligos 1, 2, and 3 (Supplementary Table 2) were mixed in DNA annealing buffer (10 mM Tris pH 7.5, 50 mM NaCl, 1mM EDTA). The mixture is heated to 95 °C for 5 min and allowed to cool down to reach RT. 54 μl of annealed dsDNA (33 μM), 90 μl of HpaII methyltransferase (222 μM), 27 μl CutSmart buffer (NEB), 27 μl SAM (20 mM), 72 μl ddH2O were mixed together to start the labelling reaction. Mixture was incubated at 30 °C for 1 h and transferred to 4 °C overnight. 500 μl of Q Sepharose beads (Cytiva) were loaded into a centrifugal spin filter and spun at 0.3x g for 1 min to remove excess liquid. Beads are then washed with ddH2O and washed with Buffer A (5 mM Tris pH 8.0, 5 mM β-mercaptoethanol). Labelling mixture was added to washed beads and inverted to mix. Increasing amounts of Buffer B (Tris pH 8.0, 2 M NaCl, 5 mM β-mercaptoethanol) were added in increments of 5-10 % and spun at 0.3x g for 1-2 min to remove excess liquid. Eluted fractions were collected and separated by electrophoresis on NuPAGE 4-12% Bis-Tris mini protein gels (ThermoFisher Scientific, NP0321BOX) and stained with Coomassie Blue (Protein Ark, GEN-QC-STAIN-1L). Fractions that contained dual-labelled DNA were pooled and concentrated. Concentrated mixture of roadblocked DNA was digested by addition of 1000 units of Exonuclease III (NEB M0206L) and 100 units T7 exonuclease (NEB, M0263L) for 1 h at 30 °C.

#### Purification of dsDNA-RecA-HpaII mini-filaments

Roadblocked DNA was mixed with RecA protein (NEB, M0249L) in 1:15 ratio (1.25x excess of RecA) in filament formation buffer (25 mM Tris-HCl pH 7.5, 10 mM magnesium-acetate, 100 mM NaCl, 1 mM TCEP, 1mM ATPγS). Final concentrations in the reaction mixture were 2 μM labelled DNA and 30 μM RecA. The filament formation reaction was allowed to proceed for 1.5 h at 37°C. 500 μl of StrepTactin Sepharose beads (IBA, 2-1201-002) were loaded into a centrifugal spin filter and spun at 0.3x g for 1 min to remove excess liquid. Beads are then washed with ddH2O, followed by a RecA wash buffer (25 mM Tris-HCl pH 7.5, 10 mM magnesium-acetate, 100 mM NaCl, 1 mM TCEP). Filament growth reaction was then added to the beads and inverted to mix. Beads were washed 9x with filament formation buffer and spun at 0.3G for 1-2 min to remove excess liquid, ensuring that ATPγS concentration was always 1mM around the beads. RecA mini-filaments were eluted with 300 μl elution buffer (25 mM Tris-HCl pH 7.5, 10 mM magnesium-acetate, 100 mM NaCl, 1 mM TCEP, 1mM ATPγS, 10 mM desthiobiotin) and spun at 0.4x g for 5 min until the beads appeared dry. The concentration of the eluted protein was determined using Bradford protein assay (Bio-Rad). The protein was then concentrated to 6 mg/ml using an Amicon Ultra-0.5 Centrifugal filter unit with 100 kDa MWCO (Millipore, UFC510024).

#### Cryo-EM grid preparation to assess RecA mini-filaments

AuFlat grids (GF-1.2/1.3-3Au-45nm-50, Protochips) were glow discharged for 40 s at 40 mA in a K100X glow discharger (EMS). For vitrification a Vitrobot^TM^ Mark IV (ThermoFisher Scientific) was used at room temperature and 100 % humidity. 4 µl of RecA mini-filaments (diluted to 1 mg/ml) was pipetted onto the grid, blotted for 11 s, and plunge-frozen without delay. Data collected at a magnification 92,000x (1.61 Å per pixel) at 200keV. The electron dose was 60 e/Å^2^. Data processing done in RELION 4.0^60^ similarly to time-resolved datasets presented below.

### Time-resolved studies of RecA homology search

#### Time-resolved cryo-EM grid preparation

Gold standard grid preparation parameters are as follows: AuFlat grids (GF-1.2/1.3-3Au-45nm-50, Protochips) were plasma treated for 1 min at 40 W power using 3 sccm of oxygen and 10 sccm of argon gas with a Tergeo-EM plasma cleaner (Labtech) using constant plasma (no pulsing). Grids were used within 20 min of plasma treatment. Grids were held at 8 mm from the nozzle. Sample was sprayed at 666 μl/min of sample flow and 4000 mg/min nitrogen gas. 0.1 CMC of OG detergent was added to protein samples immediately before loading. Variable RPM plunging was employed as described in results. When aiming for time-resolutions above 200 ms evaporative thinning time is recommended for complexes which benefit from ice significantly thinner than 60 nm. All parameters subject to change depending on grid type and biological sample.

#### Cryo-EM single-particle analysis of time-resolved RecA mini-filaments

Cryo-EM data for single-particle analysis of time-resolved RecA mini-filaments were acquired on a Titan Krios operated at 300 keV with a Falcon4i detector in counting mode with 1566 Electron Event Representation (EER) frames fractionated into 54 frames and a nominal magnification of 130,000x (0.955 Å per pixel). The electron dose was 34.5 e/Å^2^. Data pre-processing was performed in RELION 4.0 and 5.0^60^. Movies were motion-corrected by using MotionCor2 with 5 x 5 patches^61^. CTF estimation was done by CtfFind4^62^ using saved power spectra from the motion correction. Particles were picked in Topaz, with an initial hand-picked set of particles used to train the model^63^. After one round of 2D classification, the best classes were used to retrain the Topaz model for final picking. Particles were 2D classified in RELION using ‘ignore CTFs until first peak’ turned on and a regularisation parameter T=10, using the VDAM algorithm with 300 mini-batches. Sampling parameters were set to 30 pixel offset search range and offset search step of 3 pixels. 3D classification and refinement performed in RELION. Iterative rounds of Bayesian polishing and CTF refinement were performed in RELION until resolution stopped improving. Final 3D classification to identify classes was performed with a local mask containing the putative dsDNA binding regions of the filament, without alignments and regularisation parameter of T=10.

#### Model fitting and colouring

ChimeraX 1.8^64^ was used to dock PDB models. Segmentation was performed using the colour zone functionality in ChimeraX. Each map was docked separately, and certain areas were coloured manually according to interpretation.

## Acknowledgements

We thank Enchev lab members for discussions and X. Zhang and S. West for critical reading, discussions and advice. We thank the Making Lab (G. Konstantinou and W. Guan) and Structural Biology STP (A. Nans) at the Francis Crick Institute for facility support. We thank J. Brock for illustrations. Work in the Enchev laboratory was supported by the Francis Crick Institute, with funding from Cancer Research UK (CC2059), the UK Medical Research Council (CC2059), and the Wellcome Trust (CC2059) and The Crick Chris Banton Translation Fund.

## Author contributions

M.-E.M and R.I.E conceived the research. M.-E.M., J.R.M, M.M.N.E, J.A.C. and A.A. performed experiments with supervision from A.I. and R.I.E. M.-E.M, R.I.E, S.T, and M.M.N.E designed and constructed the Chronobot system. M.-E.M. and R.I.E. conceived and implemented ice thickness estimation workflows. M.-E.M and R.I.E wrote the manuscript with contributions and input from all authors.

## Data and Code Availability

Cryo-EM structural data will be deposited in the Electron Microscopy Data Bank (EMDB) to be released upon article publication. All code and list of system components to be released upon article publication.

## Ethics declarations

The Francis Crick Institute has filed three patents related to the above work and M.-E.M., M.M.N.E. and R.I.E. are named inventors. The authors declare no known additional conflicts of interest.

