## Extended Figures for "Chronobot: Deep learning guided time-resolved cryo-EM captures molecular choreography of RecA in homology search"

This document contains:

Extended figures 1-8

Extended references

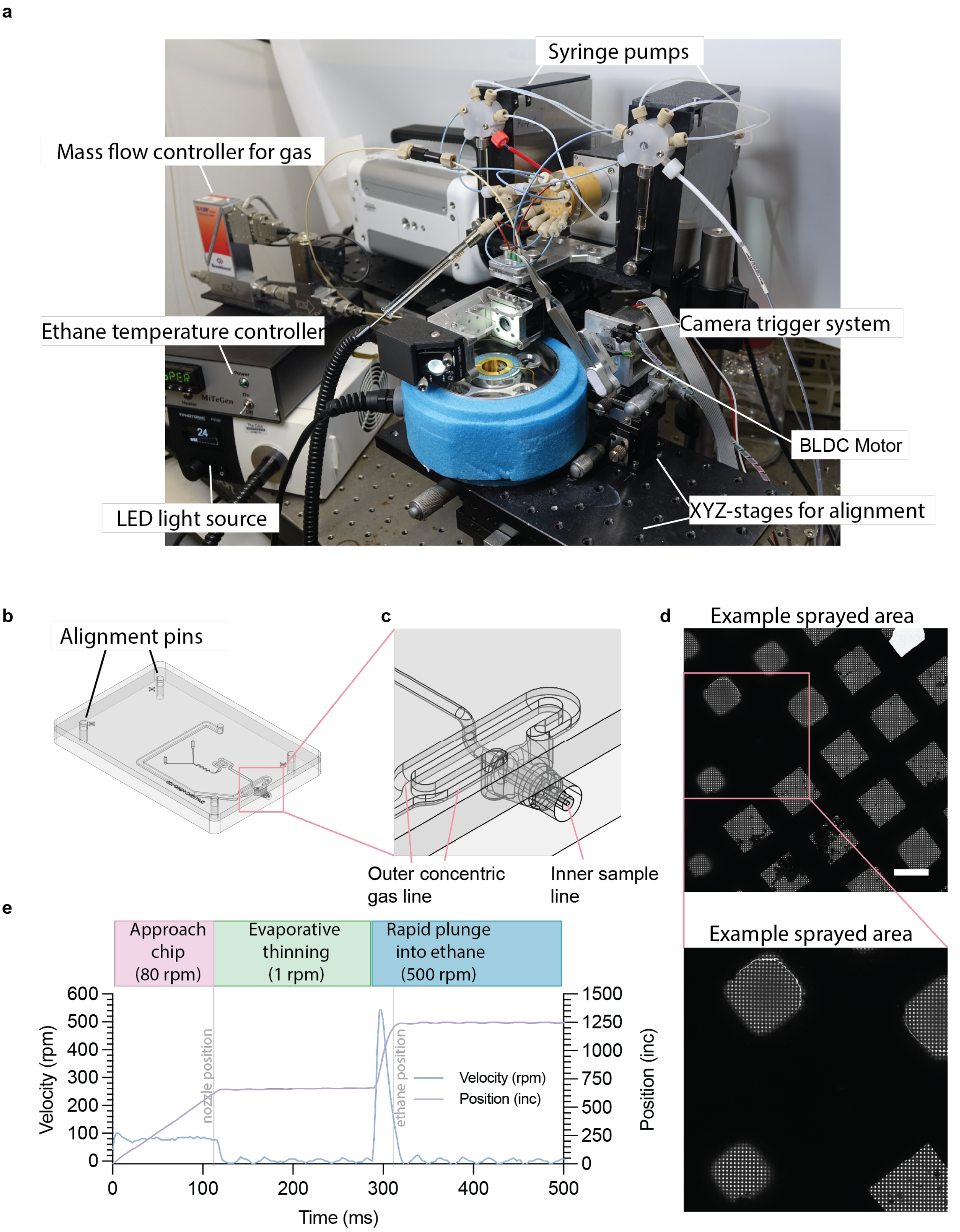


**Extended Figure 1. Mechanical setup and experimental example. a**, Photo of Chronobot with labelled system components. **b**, 3D model of a microfluidic device. **c**, inset of of b, showing a magnified view of the integrated nozzle. **d**, Representative area on sprayed grid. Scale bar = 50 μM **e**, Graphical representation of multi-rpm plunging. Position and velocity of the tweezers are shown over the course of a single plunge. Positions of nozzle and liquid ethane are shown. Position unit is internal increments of the motor. Grid is imaged during evaporative thinning. **f**, Inset of d, showing a magnified area of the micrograph.

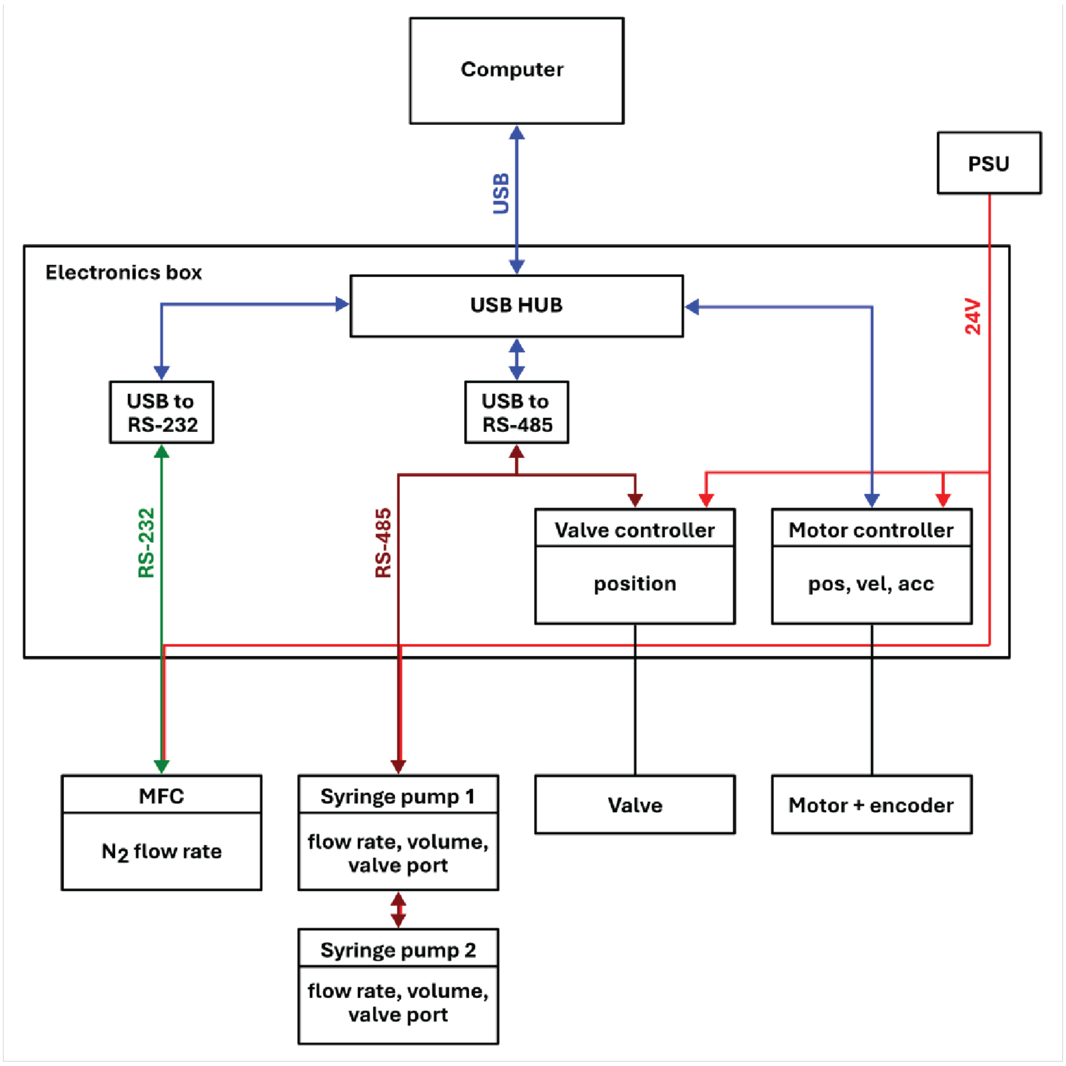


**Extended Figure 2. Electronic control diagram of the Chronobot.** Signal types depicted in different colours. Boxes underneath each component list controllable parameters.

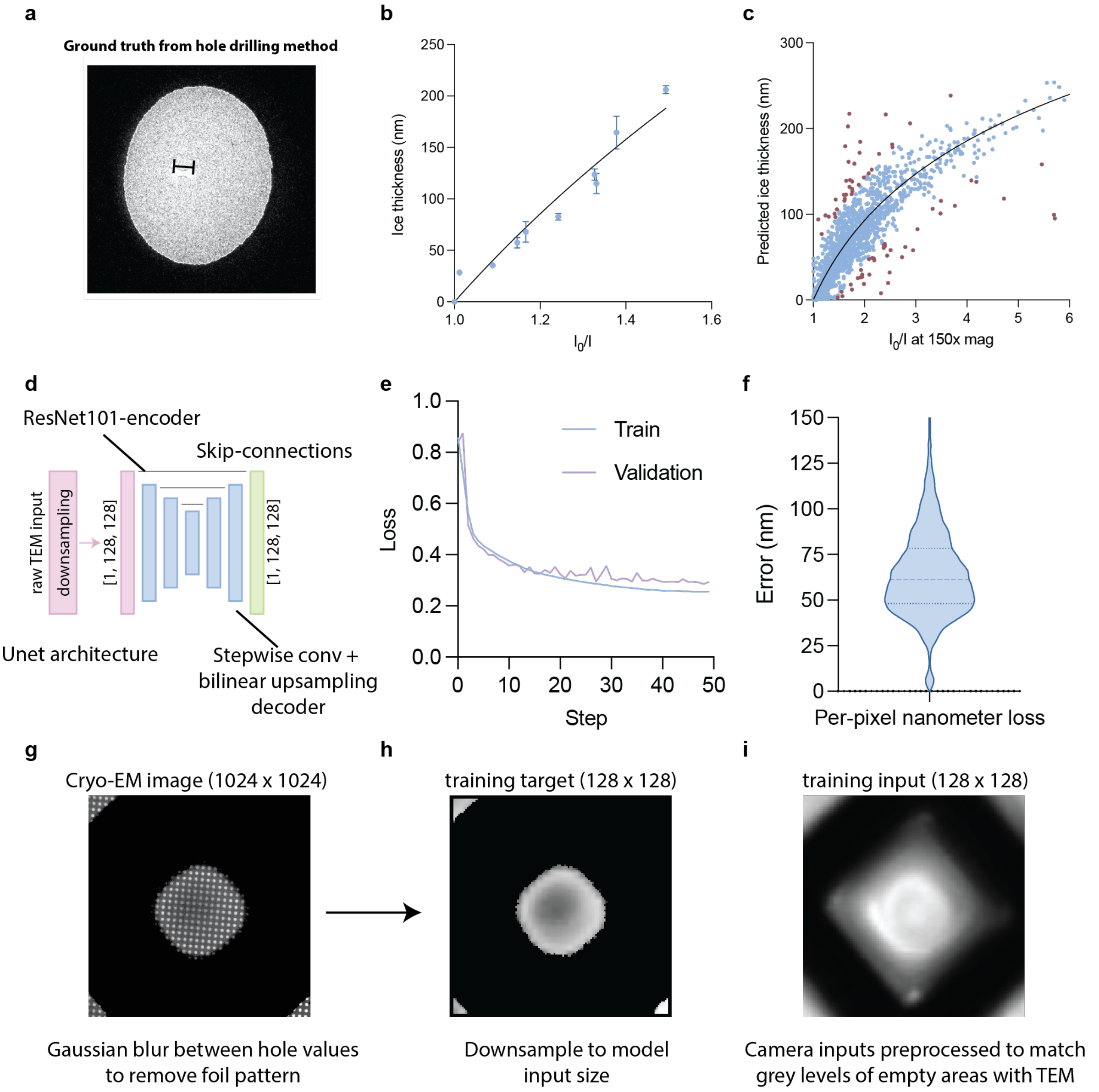


**Extended Figure 3. Ice thickness determination methods. a**, Representative micrograph of ground truth measurements collected using the ice channel method for determining ice thickness[1]. **b**, Fit of aperture limited scattering (ALS) coefficient using ice channel thickness data. **c**, Fit of ALS coefficient at 150x magnification using ice thicknesses determined using pairs of micrographs at 150 000x magnification and 150x magnification. Dots shown in red were considered outliers and not used to fit the model. **d**, Diagram of deep learning model features. **e**, Loss curves of the deep learning model during training. **f**, per-pixel evaluation of model accuracy. **g**, Image of raw input to model preprocessing. Length of the image 102 µm **h**, Representative image of preprocessed TEM image used to train deep learning model. Preprocessing smoothens the input image to remove holey foil pattern. **i**, Example of training input image, extracted from high-speed camera.

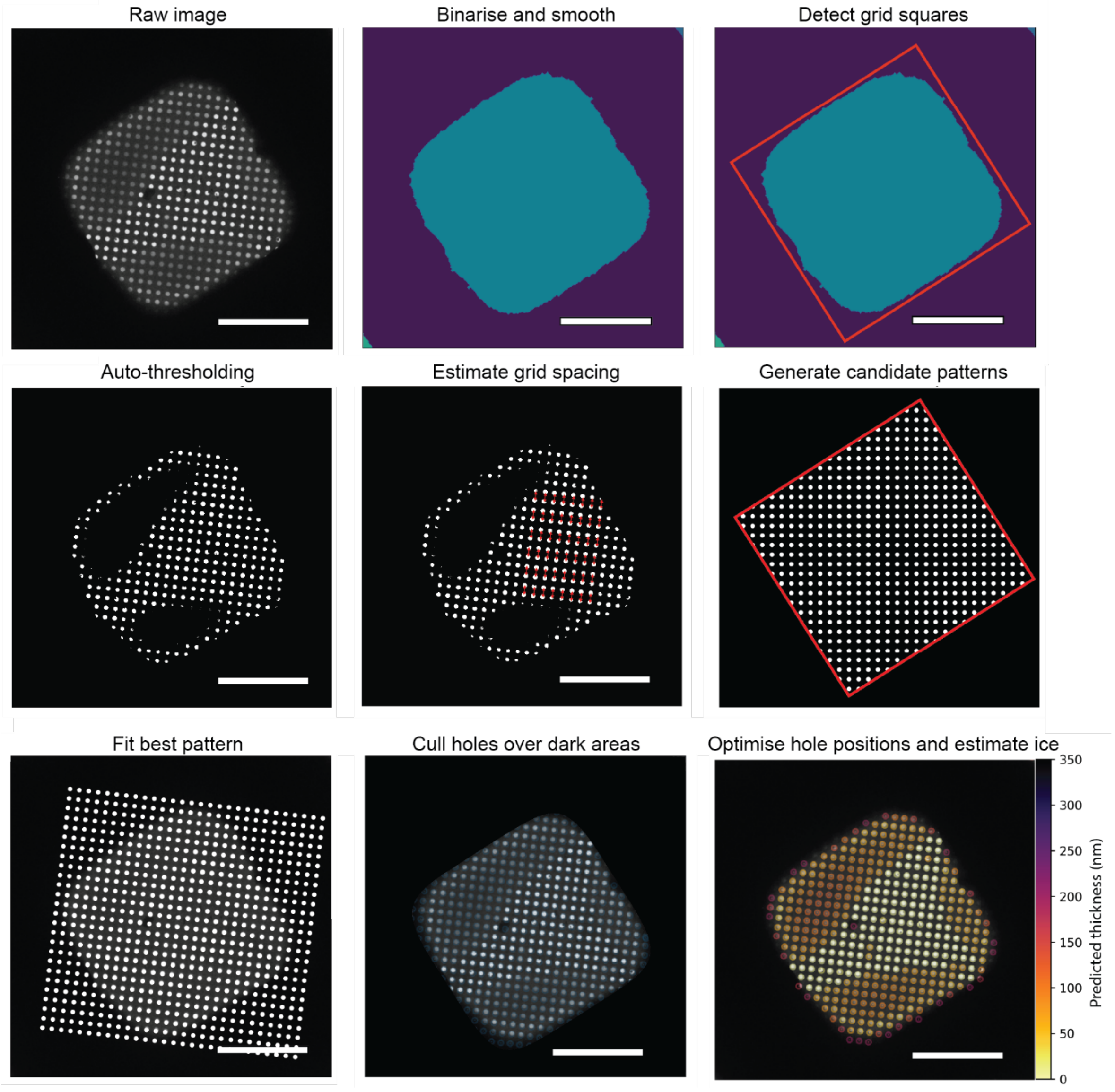


**Extended Figure 4. Algorithm for hole detection. a,** Raw image, scale bar = 25 μM **b,** Binarisation of the raw image is followed by symmetrical binary dilation and erosion steps in order to fill in the holes of the TEM sample grid gold foil. This produces a continuous representation of the square area. **c**, Squares are detected using minimal area bounding box detection in the OpenCV library. **d**, Auto-thresholding is performed using Otsu's method[2], with a fallback to manually finetuned value if fewer than 100 holes are detected on an atlas tile. **e,** Holes are detected by calculating centre of mass for each binary area in the image to obtain hole centres. Grid spacing is estimated by calculating distance and angle of 100 randomly selected points to the nearest other point. **f**, Candidate patterns are generated around +/- 1 standard deviation of the detected value from previous step. Generated patterns are culled to maintain shape of detected squares. **g**, Patterns are fit by maximising cross-correlation in Fourier space. **h**, Holes that overlay areas above a threshold value are culled as being over grid bars or ice so thick that it appears black in TEM. **i**, Finally, gradient descent is applied to all predicted hole positions to move them to the local minima around their position. To do this, the TEM image is first blurred using a Gaussian kernel with the same size as the foil holes, resulting in the centres of the holes containing the minimal values. Once positions are optimised, ice thickness values are written out.

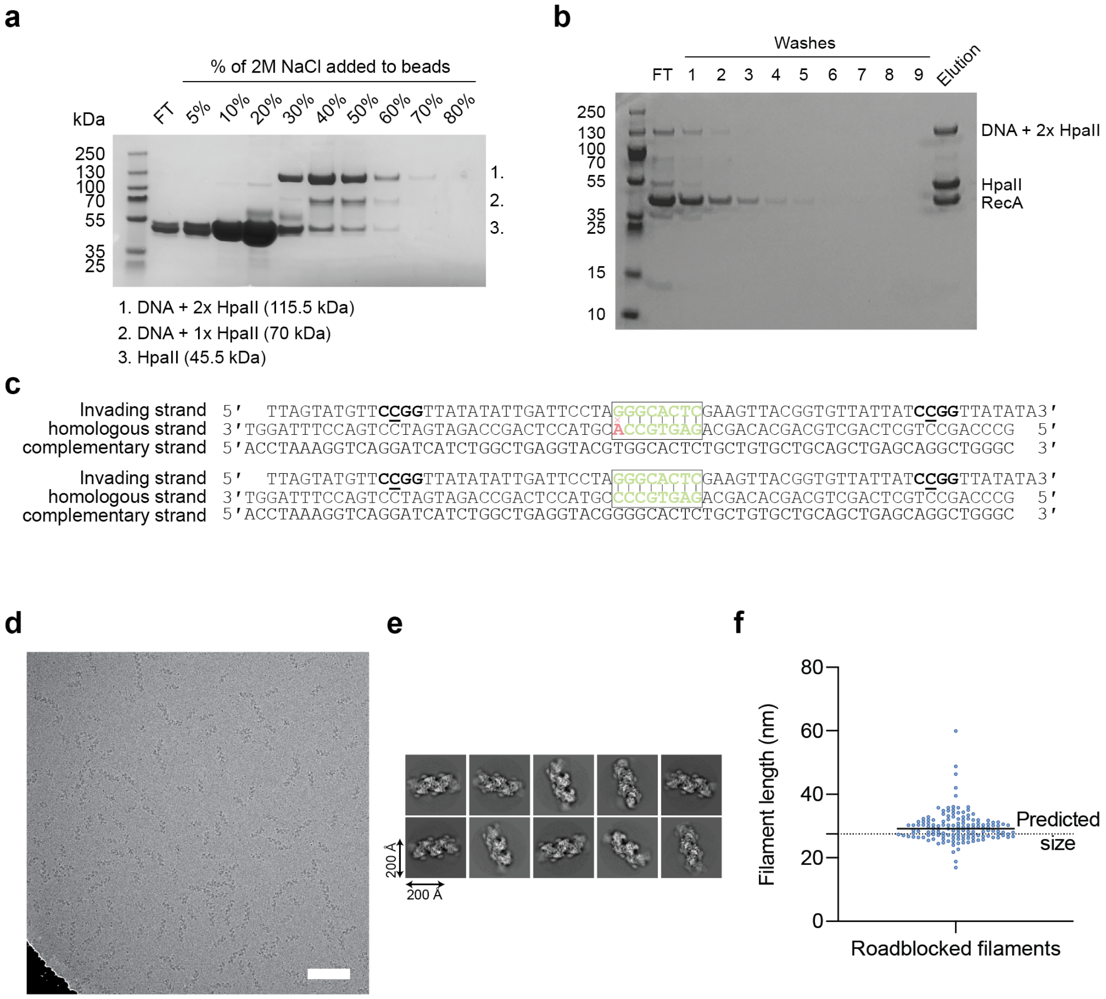


**Extended Figure 5. Preparation of roadblocked RecA mini-filaments. a**, Purification of roadblocked DNA substrates. Coomassie-stained SDS PAGE **b**, Purification of fully formed mini-filaments. Coomassie-stained SDS PAGE. **c**, Oligonucleotide designs intended for trEM studies of RecA-mediated initiation of homologous recombination. Top – design of the 7-nt substrate. Bottom – design of the 8-nt substrate. HpaII methyltransferases are covalently attached to the middle C (underlined) in the CCGG motifs on in the invading strand. **d**, Representative micrograph of roadblocked RecA mini-filaments, scale bar is 100 nm. **e**, Representative 2D classes of RecA mini-filaments, scale bars 200 Å. **f**, Manual validation of filament lengths. Measurements done in FIJI by drawing lines along visible filaments in micrographs. Predicted size is 27.5 nm. Actual measured size is higher, likely due to defocused images being used for measurement.

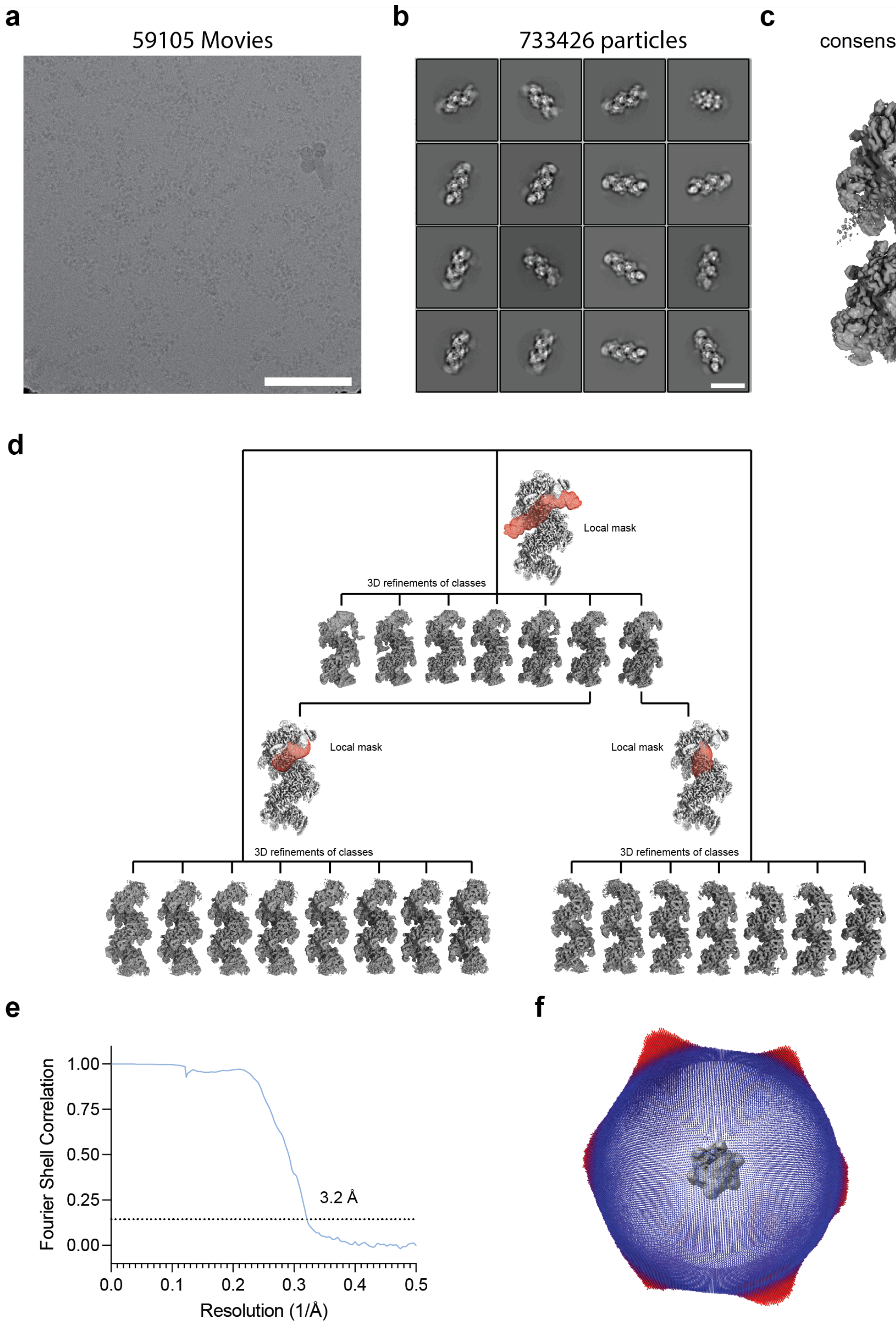


**Extended Figure 6. Cryo-EM and single particle analysis of the (RecA$^{mini\text{-}filament}$+dsDNA$^{8\text{-}nt}$)$^{250ms}$ dataset. a,** Micrograph from (RecA$^{mini\text{-}filament}$+dsDNA$^{8\text{-}nt}$)$^{250ms}$ Scale bar represents 100 nm. **b**, Representative 2D classes of the particles before consensus refinement. Scale bar represents 20 nm. **c**, Consensus reconstruction of (RecA$^{mini\text{-}filament}$+dsDNA$^{8\text{-}nt}$)$^{250ms}$. **d**, Local 3D classifications of the consensus refinement. Shown in red are local masks used for various 3D classifications. Shown classes are 3D refinement maps of each local class. **e**, Chart shows the gold-standard FSC plot of the consensus reconstruction. Dotted line marks the FSC cutoff of 0.143[3]. **f**, Angular view distribution of the consensus reconstruction.

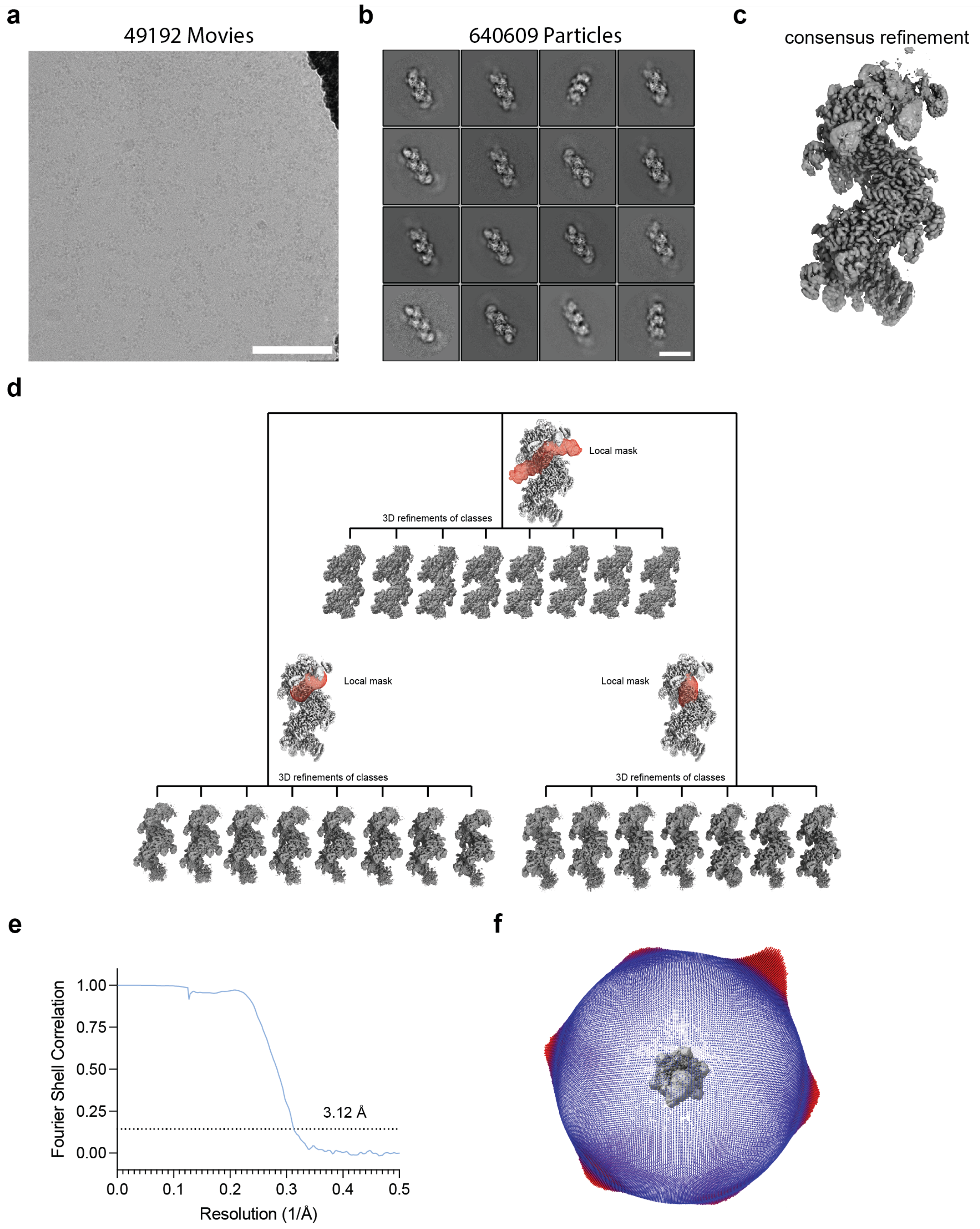


**Extended Figure 7. Cryo-EM and single particle analysis of the (RecA$^{\text{mini-filament}}$+dsDNA$^{\text{7-nt}}$)$^{\text{250ms}}$ dataset. a,** Micrograph from (RecA$^{\text{mini-filament}}$+dsDNA$^{\text{7-nt}}$)$^{\text{250ms}}$. Scale bar represents 100 nm. **b**, Representative 2D classes of the particles before consensus refinement. **c**, Consensus reconstruction of (RecA$^{\text{mini-filament}}$+dsDNA$^{\text{7-nt}}$)$^{\text{250ms}}$. Scale bar represents 100 nm. **d**, Local 3D classifications of the consensus refinement. Shown in red are local masks used for various 3D classifications. Shown classes are 3D refinement maps of each local class. **e**, Chart shows the gold-standard FSC plot of the consensus reconstruction. Dotted line marks the FSC cutoff of 0.143. **f**, Angular view distribution of the consensus reconstruction.

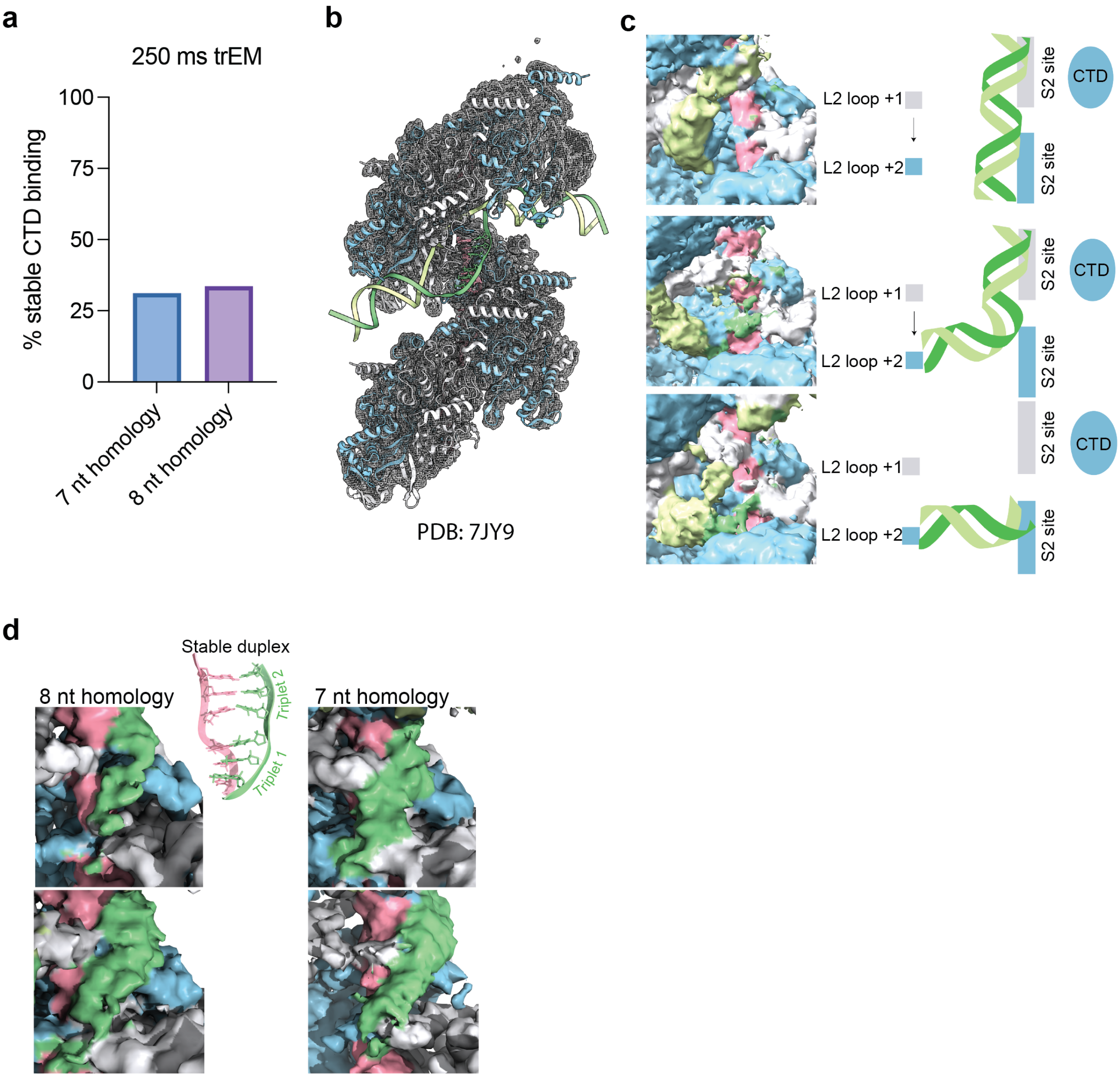


**Extended Figure 8. Structural observations of homology search intermediates. a,** Quantification of $RecA^{CTD}$ binding in both datasets. All positions of CTD binding are combined. Stable binding defined as strong density with diameter comparable to a dsDNA. **b,** Segmentation model used for all colouring (PDB-7JY9[4]) docked into the consensus refinement map of $(RecA^{mini\text{-}filament}+dsDNA^{7\text{-}nt})^{250ms}$ which is shown as grey mesh (Extended Fig 7b). Colouring model was rigidly docked into each class using ChimeraX fitmap functionality[5] . Additional densities not modelled in PDB-7JY9 were coloured manually. White and light blue represent RecA monomers. Green colours represent the homologous strand and complementary strand in a D-loop, and pink represents the invading strand. **c**, Site II binding classes as identified in both datasets. Possible interpretation illustrated to the right of each map. **d**, Comparison of homology binding densities. Compared are two classes with strongest density for homology binding in each initial 3D classification.

**Extended references**
